# Mathematically and biologically consistent framework for presence-absence pairwise indices

**DOI:** 10.1101/2021.07.14.452244

**Authors:** Arnošt L. Šizling, Petr Keil, Even Tjørve, Kathleen M.C. Tjørve, Jakub D. Žárský, David Storch

## Abstract

A large number of indices for presence-absence data that compare two assemblages have been proposed or reinvented. Interpretation of these indices varies across the literature, despite efforts for clarification and unification. Most effort has focused on the mathematics behind the indices, their relationships with diversity, and with each other. At the same time, the following issues have been largely overlooked: (i) requirement that a small re-arrangement of assemblages should only cause a small change in an index, (ii) inferences from the indices about diversity patterns, (iii) inter-dependence of indices based on their information value, (iv) overlap of the ecological phenomena that the indices aim to capture, and (v) incomparability of measures of different phenomena. Neglecting these issues has resulted in the invention or reinvention of indices without increasing their information value, although this value is crucial for correct interpretation of the indices. We offer a framework for pairwise diversity indices that accounts for these issues. We differentiate between statistical and information dependence of indices and show mathematical links between all indices, even those that have not yet been developed. Using linear algebra, we show (1) which set of indices carries complete information on assemblage arrangement, (2) how to calculate any index from two presence-absence indices, which can be used to standardize and compare different indices across the literature, and (3) what can be inferred about diversity phenomena from different informationally independent indices. It is impossible to purify an index of a single biodiversity phenomenon from the effects of other phenomena, because these phenomena inevitably constrain each other. Consequently, many recently proposed indices do not measure the phenomena that they were intended to measure. In contrast, a proper inference can be made by combining classical indices from different, information independent families.

## Introduction

Considerable effort has been made to capture variation in biological diversity using a single variable (e.g., Brualdi & Sanderson, 1999; Gotelli, 2000; Vellend, 2001; Koleff & Gaston, 2002; Gaston et al., 2007; Jost, 2007; Jurasinski et al., 2009; Tuomisto, 2010b; Carvalho et al., 2012; Chao et al., 2012; Newbold et al., 2016; Matthews et al., 2019). Differences in species composition between assemblages in space and time have been assessed by various indices, with a heated debate about which are better for different purposes (Baselga, 2010, 2012; Podani & Schmera, 2011; Schmera et al., 2020; Almeida-Neto et al., 2012; Ulrich et al., 2009, 2017). Substantial work has been done on abundance-based indices, and their inclusion in the framework of Hill numbers (Hill, 1973; Jost, 2007; Tuomisto, 2010b; Chao et al., 2012, 2014a, 2014b; Chiu et al., 2014; Barwell et al., 2015). However, accurate data on abundances are still more difficult to obtain and less common than simple presences and absences (Keil et al., 2021), particularly at large “macroecological” spatial scales. Furthermore, abundance-based indices downweight rare species, which are often of conservation interest. Because of this, pairwise indices based on presences and absences, rather than abundances, continue to dominate most global analyses of biodiversity (e.g., Dornelas et al., 2014; Blowes et al., 2019; Xu et al., 2023). The aim of this study is to provide a consistent framework for using incidence-based indices.

A particularly intensive debate has concerned the links between indices of species turnover and nestedness, and how to purify them from each other and from species richness differences. This debate can be traced back to Simpson (1943), whose invention of the Simpson similarity (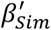 Table 1) resolved the undesirable interdependence between the Jaccard index (*J*; Jaccard, 1912; Table 1) and the difference in species richness of assemblages. Since then, a number of indices have been introduced that carry equal information to Jaccard and Simpson indices (e.g., *β*_*Sør*_; Sørensen, 1948; and *β*_*itu*_; Baselga, 2012; Table 1), or which are mathematically equivalent to earlier indices (e.g., *β*_*t*2_ = *β*_*SR*_ = *β*_*HK*_; defined by Wilson & Shmida, 1984; Schluter & Ricklefs, 1993; Harte & Kinzig, 1997; respectively; Table 1; Table R1 in Šizling et al., 2023). Yet, we lack a general framework for the relationships between indices, which would apply for any index already invented, or introduced in the future.

**TABLE 1.**
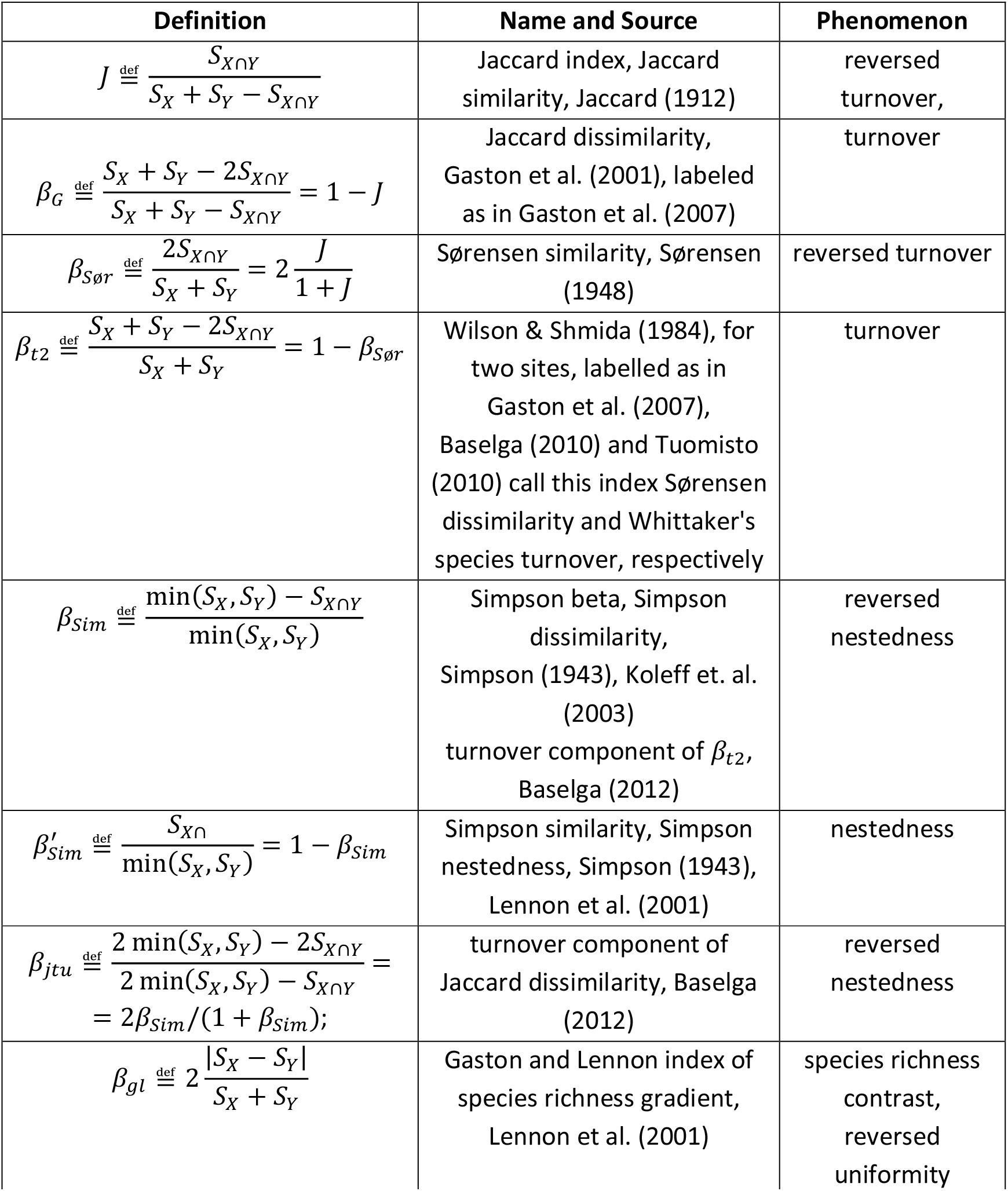
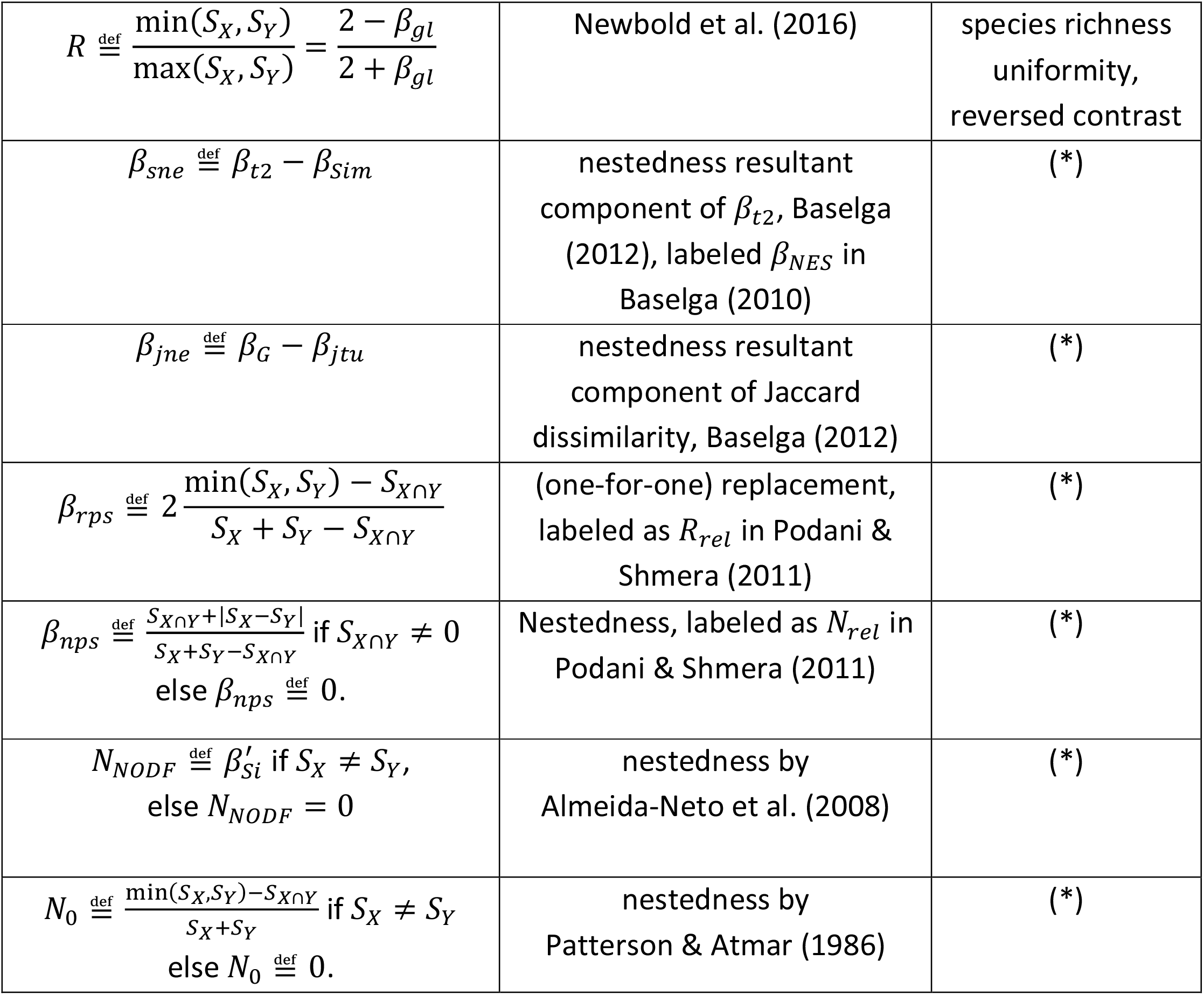
The frequently used pairwise indices. *S*_*X*_ and *S*_*Y*_ are species richness of the assemblages to compare, and *S*_*X*∩*Y*_ is the number of shared species between them. For index definitions using the a,b,c notation (*a* = *S*_*X*∩*Y*_, *b* = *S*_*X*_ − *a*; *c* = *S*_*Y*_ − *a*) see Table R1 in Šizling et al. (2023). The labelling follows Gaston et al. (2007) and the symbol 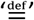 represents definitions. For a complete list of indices and their mutual relationships see Table R1 in Šizling et al. (2023). Asterix (*) labels indices that capture none of the defined phenomena.

Despite disagreements and redundancies, there has been progress. It is to Baselga’s (2010) credit that the field has refocused from the classification of indices to their meaning. Lennon et al. (2001), Tuomisto 2010b and Newbold et al. (2016) showed that the Simpson index is in fact Sørensen similarity relativized to the contrast in species richness. Koleff et al. (2003) and Legendre and De Cáceres (2013) pointed out that Simpson and Jaccard represent two different groups of indices, which were labelled as broad-sense and narrow-sense turnover. This classification resonates with Almeida-Neto et al. (2008) and Šizling et al. (2009) who realized that the Simpson similarity (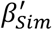; Table 1) quantifies the phenomenon of nestedness, while the Jaccard index quantifies reversed species turnover. Lastly and importantly, Ulrich et al. (2017, 2018) pioneered null models that disentangle mathematical and biological drivers of the dependence between various indices and species richness.

The realization that pairwise indices are simultaneously affected by different phenomena (e.g., by proportion of shared species, as well as, species richness difference) has recently led to an effort to purify one effect from another by means of additive partitioning. Baselga (2010, 2012), Podani and Schmera (2011) and Schmera et al. (2020) presented two ways to mathematically partition the indices. Baselga (2012) based his reasoning on the observation that some arrangements of assemblages affect both turnover and nestedness simultaneously, and the respective indices are thus ‘related’ (*sensu* Chao et al., 2012), which means that their random samples are statistically dependent (Box 1). To address this dependence, Baselga (2012) defined the “nestedness-resultant component” (*β*_*sne*_), which is 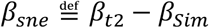, where *β*_*t*2_ is Sørensen dissimilarity and *β*_*Sim*_ is Simpson dissimilarity (see Table 1 for definitions). The subtraction supposedly removes the effect of the turnover component represented by *β*_*Sim*_ (Baselga’s 2012 interpretation) from the overall dissimilarity represented by *β*_*t*2_ (Baselga’s 2012 interpretation), and what remains is considered to be the ‘nestedness resultant component’. In contrast, Podani and Schmera (2011) see nestedness as antithetic to replacement (their synonym for turnover) and so indices of nestedness (*β*_*nps*_; Podani & Schmera, 2011) and replacement (*β*_*rps*_; Podani & Schmera, 2011) (see Table 1 for definitions) together sum to one. Their index of nestedness is thus 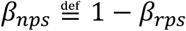.

We argue that the abovementioned approaches have several problems (outlined in the section below) which has led to a situation where researchers are typically not sure which indices can be used in different occasions and what the indices actually measure. To sort out the confusion, we present a unified framework for incidence-based indices that addresses these issues (see also Tuomisto 2010a, 2010b). In our framework, the indices are tools to capture phenomena observed during fieldwork or data processing, and we show that these phenomena are already mutually dependent by their very nature, regardless of the indices that quantify them. This makes statistically independent or unrelated indices impossible. We further show that the interdependence between the phenomena makes it impossible to ‘purify’ any index from the effects of other phenomena.

Instead of statistical independence (hereafter *s-independence*) or unrelatedness (sensu Chao et al., 2012) of indices, we advocate for accounting for the independence of the indices in terms of their information content, which we call here *i-independence* (Box 1; note that in mathematics this is called simply independence). We then show that only two i-independent indices, together with species richness, carry enough information to compute any (even not yet introduced) index, and we provide a user friendly tool to perform this conversion. We also develop guidelines to attribute the indices to different diversity phenomena. We demonstrate that particular types of interrelation of assemblages introduce mutual i-dependence between otherwise i-independent indices, and that this effect can obscure differences between otherwise clearly distinguishable patterns. Finally, we show that all the mathematically and ecologically meaningful indices have already been described by Jaccard (1912), Simpson (1943) and others in the early 20^th^ century (see also Tjørve et al., 2018).

### Problems with previously suggested indices

We distinguish five fundamental problems with the indices proposed in the literature during the last few decades. Our basic assumption is that different indices should measure different – albeit not necessarily entirely independent – phenomena, and that the value of an index should reflect the strength of a given phenomenon (Ulrich et al., 2017, 2018). The problems listed below represent cases when this general assumption is not fulfilled.

#### Problem 1

Some commonly used indices do not satisfy the *requirement of continuity* (Neumann & Morgenstern, 1953). This requirement, presented for ecologists by Podani and Schmera (2012), states that a *small* re-arrangement of species assemblages should lead to only a *small* change in the index value and conversly, a *small* change in an index value should indicate only a small re-arrangement of the assemblages. This is important for two reasons: an index that changes considerably with a negligible re-arrangement of assemblages is (i) sensitive to errors in observed data, and (ii) leads to misleading inference about the pattern and/or ecological phenomenon. For example, common indices of nestedness as *N*_0_(Patterson & Atmar, 1986) and *N*_*NODF*_ (Almeida-Neto et al., 2008) have been defined as zero when species richness of one site (*S*_*X*_) is equal to species richness of the other site (*S*_*Y*_). This violates the requirement of continuity (Figure 1). To satisfy the requirement when *S*_*X*_ = *S*_*Y*_, a value of nestedness should instead fall as close as possible to the nestedness values for *S*_*X*_ = *S*_*Y*_ − 1 and *S*_*X*_ = *S*_*Y*_ + 1. Similarly, the problematic assumption of zero nestedness for *S*_*X*_ = *S*_*Y*_ also affects the “nestedness-resultant components” by Baselga (2010, 2012) (Figure 1). Baselga (2010, 2012) explicitly stated that, in the absence of nestedness, an index of dissimilarity equals its turnover component. However, this would hold only if the equality of *S*_*X*_ and *S*_*Y*_ implied zero nestedness (Figure 1). The problem is that two precisely identical assemblages should be, according to the requirement of continuity, similarily nested as two almost identical and absolutely nested assemblages, i.e. they must not have zero nestedness.

**FIGURE 1.**
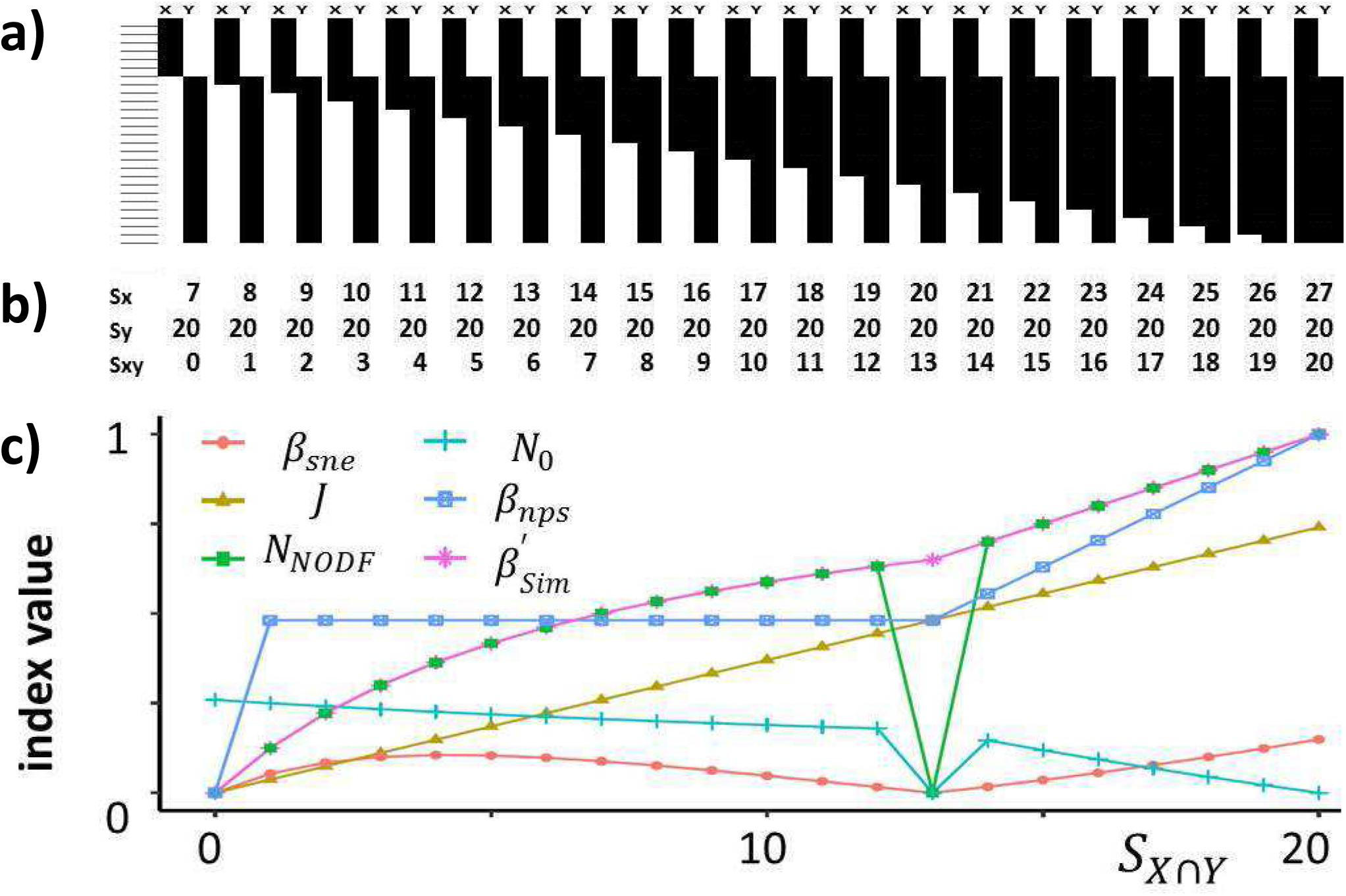
The requirement of continuity (see section *Problems with previously suggested indices*), and consequences of its violation. The top plot (a) shows a sequence of 20 pairs of assemblages X and Y, with a continuous change of the arrangement of assemblages, increasing similarity in species composition and nestedness from left to right, and constant total species richness. Columns show assemblages, lines species and blackened cells presences of species within assemblages. The first two lines in (b) show numbers of species in the left (Sx) and right (Sy) assembages for each situation along the gradient. The third line in (b) shows *S*_*X*∩*Y*_ labeled as Sxy. The most bottom plot (c) shows the values of *β*_*sne*_ (red circles), *J* (olive triangles), *N*_*NODF*_ (green squares), *N*_0_ (green-blue crosses), *β*_*nps*_ (blue squares), and 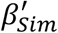 (purple asterisks).

#### Problem 2

The relationships between indices and ecological phenomena that the indices aim to capture (e.g., nestendess) have been typically derived using an ambiguous *from-phenomenon-to-index* implication: a particular state of the phenomenon (e.g., high or low nestedness) affects the value of an index, so it is assumed that the index measures the phenomenon. However, if an index is to be useful, the appropriate reasoning should follow a *from-index-to-phenomenon* implication: a value of the index should *always* correspond to a unique state of the phenomenon (e.g., only to high or only to low nestedness, not both). In other words, one index value should not be assigned to multiple different states of the phenomenon (see the problem of *converse logic*; e.g., Audi, 1999). An example of the ambigous from-phenomenon-to-index implication is used by Podani and Shmera (2011), who argue that “*because species replacement and nestedness reflect contrasting ecological phenomena […] it is meaningful to express nestedness (β*_*nps*_*) with the effect of species replacement (β*_*rps*_*) completely removed from [its] maximum*”. However, in Figure 1 we show that cases with 1-13 shared species have equal *β*_*nps*_, although they apparently have different nestedness, conceptually predefined as the extent in which species of smaller community occur also in the larger community (Podani and Schmera, 2011; Patterson, 1984; Baselga, 2010). The equality between their *β*_*nps*_ thus violates the from-index-to-phenomenon implication. Similarly, the fact that the absence of nestedness leads to *β*_*Sør*_ = *β*_*Sim*_ (Baselga 2010, 2012) does not mean that *β*_*Sør*_ = *β*_*Sim*_ implies no nestedness. This problem of the incorrect implication was mentioned by several authors (e.g., Ulrich et al., 2017, 2018; Schmera et al., 2020) but has not been appreciated in practice.

#### Problem 3

So far, attempts to remove an effect of one index from another have accounted for *s-dependence* (statistical dependence) between indices (e.g., Simpson, 1943; Koleff et al., 2003; Baselga, 2010, 2012) but failed to account for *i-dependence*, i.e. the dependence of indices in terms of their information content (Box 1). However, in terms of the exact meaning of the indices and their links to ecological phenomena, i-dependence is more fundamental, as we will show later. In short, two indices are considered i-dependent if one can be uniquely determined by the other and vice versa (Box 1).

#### Problem 4

The mathematical operation of *subtraction* of indices does not remove one effect from another. Baselga (2010, 2012) has suggested that subtraction of an index of turnover component from an index of dissimilarity leads to the nestedness-resultant component, irrespective of the indices used. Baselga (2012) applies the reasoning about nestedness-resultant components equally to *β*_*t*_ and *β*_*G*_, so that the nestedness-resultant component of Sørensen dissimilarity (*β*_*sne*_; Table 1) is *β*_*t*_ − *β*_*Sim*_, and the nestedness-resultant component of Jaccard dissimilarity (*β*_*jne*_; Table 1) is *β*_*G*_ − *β*_*itu*_ (*β*_*Sim*_, and *β*_*itu*_ are the ‘turnover components’ of Sørensen and Jaccard dissimilarities, respectively (see Table 1). However, the subtraction can remove the effects only for *β*_*t*_ or *β*_*G*_ (or none of them), but not for both, and it is unknown which of *β*_*sne*_ and *β*_*jne*_ would be the ‘true’ nesteness resultant component. The reasoning is as follows: *β*_*t*_ is i-dependent to *β*_*G*_ (one can be calculated from the other) and, similarly, *β*_*itu*_ is i-dependent to *β*_*Sim*_ (see Table 1, and Thesis T01.21 in Šizling et al., 2023). If the subtraction produced nestedness-resultant components in both cases, then the subtraction would have to account for the same effect in both cases, and thus *β*_*sne*_ and *β*_*jne*_ should also be i-dependent. However, it is not the case (Baselga, 2012). The problem is that the minus operator in 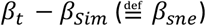 has a different meaning than the minus operator in 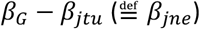; similarly the subtraction in *x* − *y* does not have the same meaning as the subtraction in *log x* − *log y* (since *log*(*x*) − *log*(*y*) = *log*(*x*/*y*)), even though *x* and *log x* are i-dependent. As a result, the *log*ic that subtraction eliminates an effect cannot be applied to both *β*_*sne*_ and *β*_*jne*_, and only one of them (if any) can capture the nestedness-resultant component, and we don’t know which one.

#### Problem 5

More generally, although pairwise indices are dimensionless quantities and can take values within the same range (e.g., 0 and 1), they are not universally comparable (ISO, 2009; Schmera & Podani, 2011). For example, the framework introduced in Baselga (2010) subtracts and compares Sørensen dissimilarity, *β*_*t*_, and Simpson dissimilarity *β*_*Sim*_ (Table 1); however, these all are not commensurable, as pointed out by Schmera and Podani (2011), who proposed to sum (or subtract) only indices with the same denominator.

### Theory

Here we first describe four distinct phenomena that represent different aspects of the structure of species communities, and can be quantified by different indices. Second, we define the constraints necessary to attribute each of the indices to one phenomenon. Third, we define the independence of the indices (including alpha diversity) in terms of their information content. We contrast this i-independence with s-independence (including relatedness sensu Chao et al., 2012; Figure 2; Box 1), which was a major topic of a large ecological literature (Simpson, 1943; Koleff & Gaston, 2002; Jost, 2007; Baselga, 2010).

**FIGURE 2.**
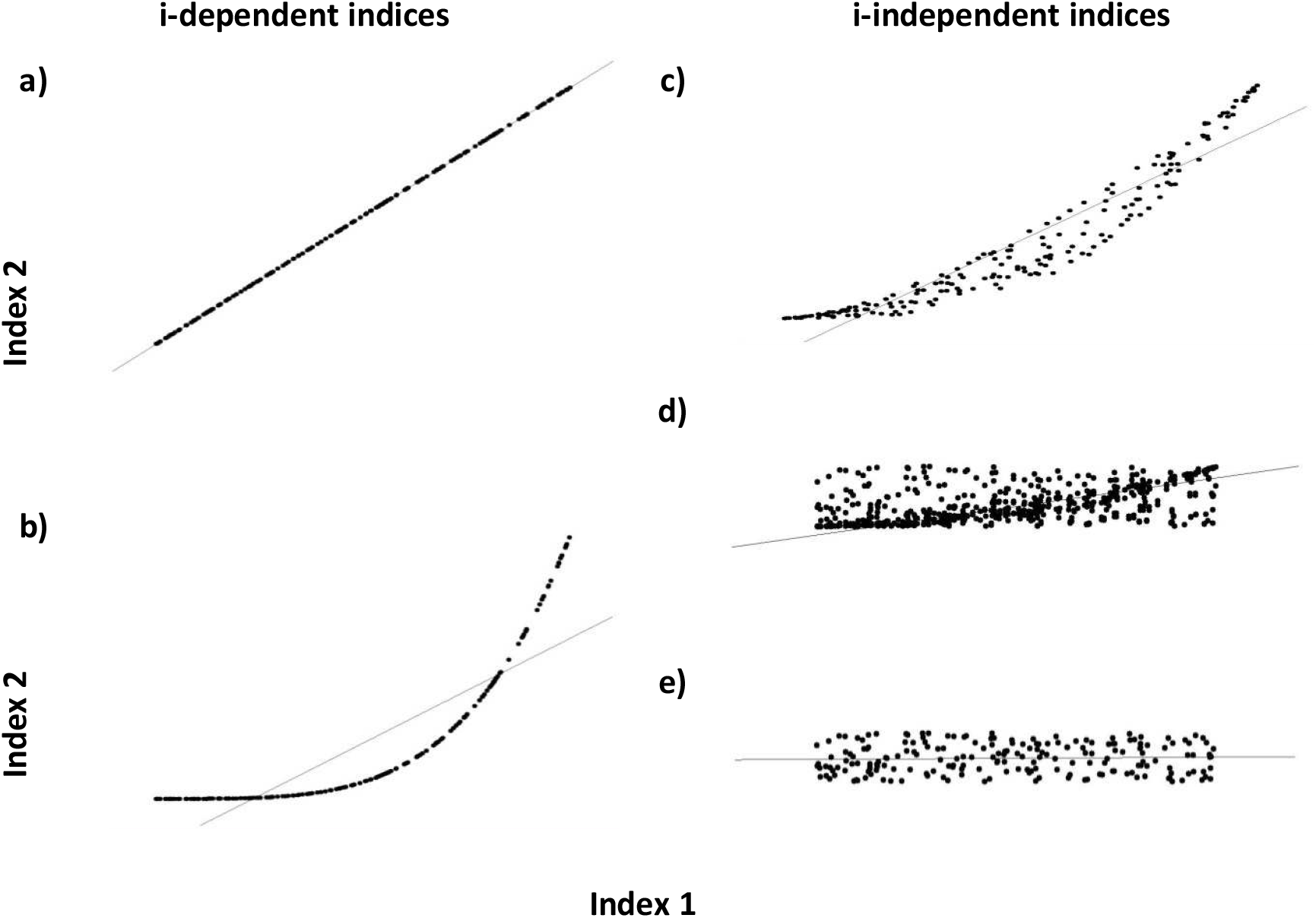
An example of relationships between two indices that are i-dependent (a,b), i-independent (c,d,e), s-dependent (a,b,c,d), s-independent (e), related sensu Chao et al. (2012) (a,b,c) and unrelated (d,e) indices. The dots show 200 samples from the underlying relationships and the lines show linear regressions. I-dependent indices (a,b) carry equal information because the strictly increasing or decreasing relationship between them uniquely transforms one index to another. Relatedness (Chao et al., 2012) is one of the mechanisms that drive s-dependence, and i-dependence is a special case of relatedness where both upper and lower constraints are identical (a,b). The existence of identical constraints must be proven by means of mathematical analysis using the formulae that define the indices for each particular case separately (Thesis T01 in Šizling et al., 2023). Anyway, there are apparently no one-to-one (strictly increasing nor decreasing) continuous lines beyond the samples of i-independent indices (c,d,e) (which does not require analytical proof). Only the indices in panel (a) scale in a linear manner. The indices in panel (b) are continuous (non-linear) transformations of each other. Indices in panels (c,d,e) can also be viewed as transformations of each other, but this transformation is not a homeomorphism and thus each index may or may not measure different phenomena, depending on the constraints (Figure 3). The degree of s-dependence measured by Pearson correlation coefficient decreases from (a) to (e) (1; 0.95; 0.86; 0.73; 0; N=200). Panels a and b show i-dependent relationships, panels c,d, and e show i-independent relationships.

#### Predefinitions of four diversity phenomena

We propose that most of the commonly used indices have originated as proxies for *four phenomena*: (1) nestedness, (2) species turnover, (3) species richness contrast, and (4) beta-diversity sensu stricto (i.e. the contrast between small-scale and large-scale richness). These four phenomena correspond to different (but strongly interrelated) *observations*, where: (i) species of the species-poor site are more or less present in the species-rich site; (ii) species lists vary between the visited sites (but the species richness may or may not change); (iii) species richness varies negligibly or considerably between visited sites, and (iv) species richness of a site represents lower or higher portion of species richness of the whole region.

Imagine an ecologist roaming the landscape, leaving one site, and approaching another, detecting changes in species richness, changes in species composition, and changes in overlap in species composition between species assemblages. They always observe a combination of these four patterns, as the phenomena are interconnected and constrain each other. Apparently, if there is no turnover, then all assemblages are identical and there is no richness contrast. The loss of a species from species poor assemblages increases both the richness contrast and turnover, and perfectly nested assemblages show lower turnover than non-overlapping assemblages of the same richness. The interdependence and mutual constraints between the phenomena are their inevitable properties, and are consequently reflected by the indices. Nevertheless, these four patterns delimit the whole range of possibilities. So, the ecologist can intuitively perceive the phenomena in the field even without the knowledge of the mathematically defined indices (similarly, we can compare the speed of a moving object without measuring it). The four phenomena are:

##### Nestedness

The idea of nestedness was originally intended to capture the patterns in assemblages of archipelagos and/or of inland islands (e.g., mountain ridges). Patterson (1984), and Patterson and Atmar (1986) noticed that species on species-poor mountain ridges were almost universally found at the species-rich sites. They suggested that this pattern was driven by selective extinction and called it a “nested pattern”. Their index of nestedness, *N*_0_, was designed to measure the deviation from a perfectly nested assemblage, in which species-poor sites have no unique species. This concept of nestedness fully agrees with the mathematical definition of nestedness where a subset is nested within the set. Although the original indices of nestedness (*N*_0_, *N*_*c*_, and Discrepancy; Patterson & Atmar, 1986; Wright & Reeves, 1992, and Brualdi & Sanderson, 1999) were designed for multisite assemblages, here we explore them using only two assemblages (Koleff et al., 2003; Gaston et al., 2007), provided that the indices for multi-site assemblages are extensions of two-site indices.

##### Species turnover and replacement

Generally, species turnover addresses the gain and the loss of species in space or time (Cody, 1975; Wilson & Shmida, 1984; Tuomisto, 2010a). Turnover is sometimes narrowed down to *replacement*, defined by Podani and Schmera (2011) and Schmera et al. (2020) as a situation where one species replaces exactly one other species, irrespective of the ecological role or habitat requirements of the species. This one-for-one replacement is equivalent to Balselga’s (2010) idea of turnover component (Schmera et al., 2020), because Baselga’s (2010) aim was to separate turnover from species richness contrast. Here we follow the most common approach (Lennon et al. 2001, Koleff et al. 2003, Gaston et al. 2007) and we understand turnover as a more general phenomenon referring to a change of species composition between assemblages.

##### Species richness contrast

Species turnover between sites, i.e. the difference in species composition (Cody, 1975; Lennon et al., 2001), can be determined by the difference in species richness (Harrison et al., 1992). Although these two differences are bound by each other (a difference in species richness between two assemblages is always accompanied by difference in their species composition), they are also mutually independent to some extent (two equally species rich assemblages can have all or no species in common). Here we adopt the idea that the contrast between the species richness of two different assemblages (called species richness gradient in Lennon et al., 2001) and species turnover are two distinct phenomena (Lennon et al., 2001) although they are not mutually exclusive - these two phenomena overlap each other and their measures must therefore correlate with each other.

##### Beta-diversity

The original idea behind beta-diversity is that different regions have different relationships between local (alpha) and regional (gamma) diversity (Whittaker, 1960). Beta-diversity thus quantifies the contrast between gamma-diversity and average alpha-diversity (Whittaker, 1960). Therefore, beta-diversity does not primarily compare two assemblages with different locations, but instead a set of sub-assemblages with a merged assemblage of the whole region. The Whittaker (1960) formula that defines beta-diversity (hereafter beta-diversity *sensu stricto*) was later included in the indices of similarity between two different assemblages (Koleff et al., 2003) and several mathematical links between similarity indices and Whitaker beta-diversity *sensu stricto* were highlighted (Koleff et al., 2003; Tuomisto, 2010b; Chao et al., 2012). However, in accord with Koleff et al. (2003), Jost (2007) and Tuomisto (2010b) we contend that the two forms of comparison (i.e., between species richness of an assemblage and its sub-assemblages, and between two separate assemblages) should not be confused.

#### Formal definitions of the phenomena based on their constraints

Here, we show the strict constraints that delimit the extreme cases of the phenomena described above. This will serve to attribute each index to a particular phenomenon, which is a prerequisite for any meaningful index. To describe the constraints mathematically, we will use six extreme re-arrangements of two sets as illustrated in the Venn diagram in Figure 3. Each set can be considered as a list of species. The constraints are:

**FIGURE 3.**
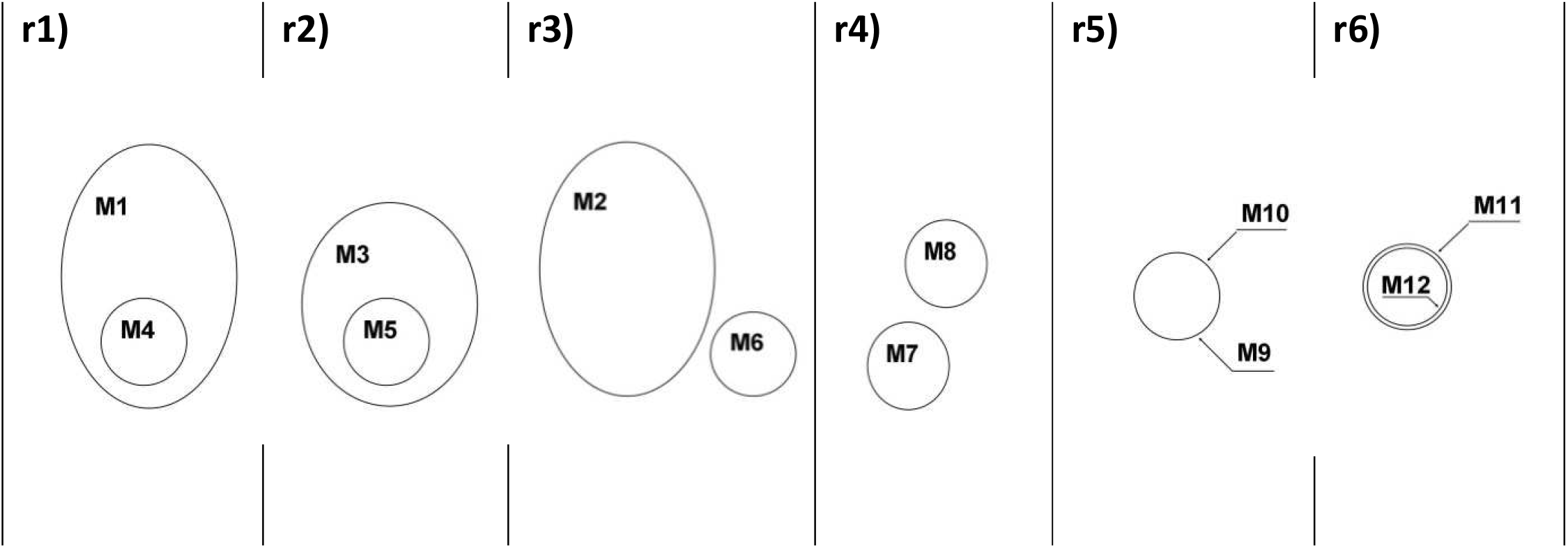
Constraints by the three (excluding beta-diversity *sensu stricto*) diversity phenomena on the indices. Various re-arrangements of two sets (lists of species) in the Venn diagrams define differences between the phenomena. The sizes of the sets follow M1 = M2 > M3 > M4 = M5 = M6 = M7 = M8 = M9 = M10 =M11>M12. The sets M9 and M10 are identical. The constraints that define the spatial phenomena are as follows: Nestedness: Min = N[r4] = N[r3] < N[r1] = N[r2] = N[r6] = N[r5] = Max; Turnover: Min = T[r5] < T[r6] < T[r2] < T[r1] < T[r4] =_(1)_ T[r3]; and Richness Contrast: Min = Ct[r5] = Ct[r4] < Ct[r6] < Ct[r2] < Ct[r1] = Ct[r3]. The ‘*Min’* and ‘*Max’* label the extreme values of the indices (usually *Min*=0 and *Max*=1) and the letters in brackets refer to the re-arrangements. The equation =_(1)_ is an arbitrary choice to separate the species richness contrast from species turnover.

- *Nestedness* has its maximum where all the species from the species-poor assemblage are found also in the species-rich assemblage, and minimum where the two assemblages have no species in common (e.g., Patterson & Atmar, 1986; Wright & Reeves, 1992; Brualdi & Sanderson, 1999; Baselga, 2010). Therefore, the two assemblages are maximally nested if one of them is completely contained within the other, regardless of their size (as in re-arrangements r1, r2, r5, r6 in Figure 3).
- *Turnover* captures the contrast in species composition between two or more assemblages. Consequently, in Figure 3, the turnover found in re-arrangement r3 must be larger than that of r1, which in turn represents a larger turnover than r2. In r5, the species lists are identical, hence there is no turnover at all, which means that the index of turnover is at its minimum. In the section above, we distinguish turnover (T) from species richness contrast ( Lennon et al., 2001) and so we state, in accord with Tuomisto (2010a), that *T*[*r*3] = *T*[*r*4] but *C*[*r*3] > *C*[*r*4] (Figure 3), i.e. if no species are shared, turnover is maximal regardless of the species richness contrast.
- Because we define the *species richness contrast* as simply the contrast in species richness between two sites, the re-arrangements r1 and r3 in Figure 3 represent the same species richness contrast, as do r4 and r5 in Figure 3. The latter also represents a minimum species richness contrast.

#### Independence of indices in terms of information content

To capture different phenomena by different indices, we need an idea of an independence of indices in terms of their information content – only those indices that carry different information can distinguish different phenomena. To find out the information provided by an index, we need to see the definitions of the indices as equations to solve (Box 2). When we add an equation to the set of *n* equations, and if this new set of equations provides an identical solution to the previous set, then the new equation is said to be *i-dependent* on the others and carries no extra information. If the solution of the *n*+1 equations is a subset of (but not equal to) the solution of the *n* equations, then the new equation is *i-independent* of the others and carries extra information. Note that this is a widely accepted mathematical definition of independence in a system of equations (see also Box 2) and that it follows the theorem from information theory that only the variables that are uniquely mapped to each other have equal information content (Orlitsky, 2003).

We utilize this mathematical definition of independent equations and define the indices that are mutually i-independent if the equations of the indices are mutually independent (Box 2). In extreme circumstances, the set of equations with one unique solution carries complete information on both assemblages. It all means that the complete set of equations uniquely determines richness of both assemblages (*S*_*X*_, *S*_*Y*_), as well as species overlap (*S*_*X*∩*Y*_; for details see Box 2).

For example, 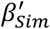 (Simpson nestedness) and *J* (Jaccard similarity) (Table 1) are i-independent because there is no way to convert 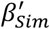 to *J*. This is easy to see from Figure 4a,b where more than one value of 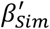 is attributed to each value of *J* (*J* ≠ 0). This is an analogy to a case from physics, where the kinetic energy (*E*_*k*_) and momentum of a moving object cannot be calculated from each other although they are computed using the same variables (*E*_*k*_ = 0.5*mv*^2^, *p* = *mv*; *m* is mass and *v* is speed). It induces relatedness (according to Chao et al., 2012; see Box 1), but *not one-to-one* correspondence (Figure 2) between energy and momentum (Figure 4c,d). However, because *E*_*k*_ and *p* are independent in terms of the lack of *one-to-one correspondence*, the equations of these variables can be combined to determine the mass and speed of an object. The indices *J* and 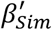 carry different information about assemblages, similarly to *E*_*k*_ and *p* that carry different information about moving objects, although statistical tests would reveal their mutual statistical dependence, and the constraints reveal their relatedness.

**FIGURE 4.**
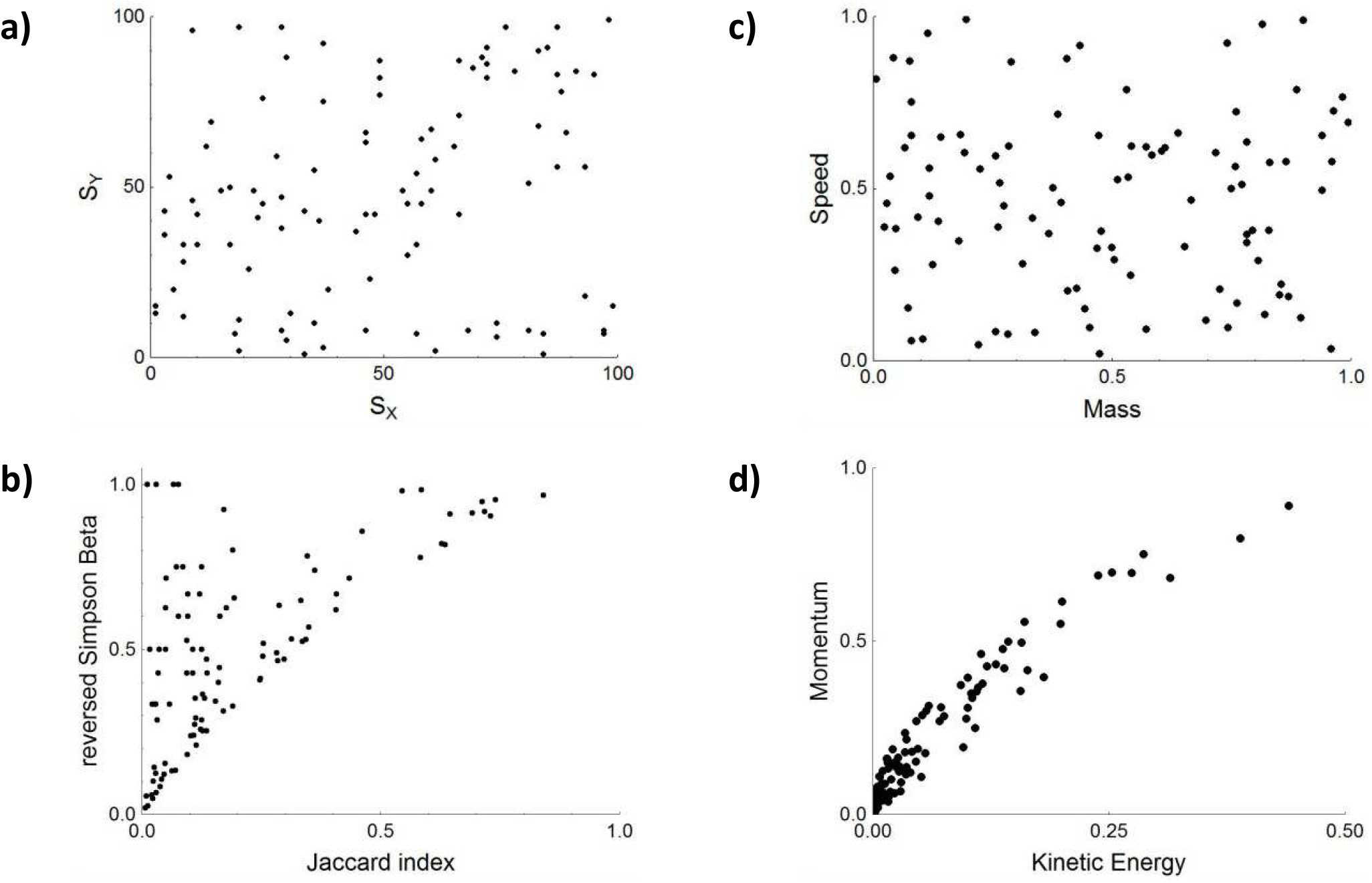
Statistical and informational (in)dependence in ecology (a,b) and physics (c,d). Mutually statistically independent (s-independent) values for species assemblages in ecology (*S*_*X*_ and *S*_*Y*_) (a), and moving objects in physics (mass and speed) (c) do not necessarily result in statistically independent indices (*J* and 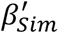) (b) and variables (*p* and *E*_*k*_) (d). This is despite the fact that 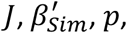 and *E*_*k*_ carry a specific information and therefore are i-independent regarding their information value. Here *S*_*X*_, *S*_*Y*_ (∈ {1, …, 100}) and *S*_*X*∩*Y*_ (≤ *min*(*S*_*X*_, *S*_*Y*_)) are drawn at random from a uniform distribution.

## Results

### Families of indices

In the previous section, we have developed a framework to assign the indices to different phenomena. Now, we will group them to families within which indices share equal information, and thus capture the same phenomena. Then we proceed by assigning a phenomenon to the index which is used most often or is more convenient for some reason. Finally, we will show how to convert indices within their respective family to each other (Box 3). Mathematical details of these steps are in repository (Theses T01-T20 in Šizling et al., 2023), and here we summarize the results.

There is a number of pairwise indices and each has some information value. Some of them, however, are i-dependent, and they are thus *equivalent* in terms of any inference. To distinguish families of i-dependent indices, we first examined their mutual bivariate relationships, i.e., we simply plotted the indices against each other, using simulated and empirical data (Figure 5). Where we did not see a one-to-one correspondence (Figure 2), i.e., the line joining the points in the plot was neither strictly increasing nor strictly decreasing, the two plotted indices were considered i-independent. Where we found a one-to-one correspondence along a strictly increasing or decreasing curve, we had to prove the i-dependence by mathematical analysis (Thesis T01 in Šizling et al., 2023). In this way, we found several families of mutually equivalent indices (Figure 5), and our mathematical analysis confirmed the one-to-one correspondences (i-dependence) between the indices within these families. The three families of more than one index can be attributed to turnover, nestedness, and species richness contrast.

**FIGURE 5.**
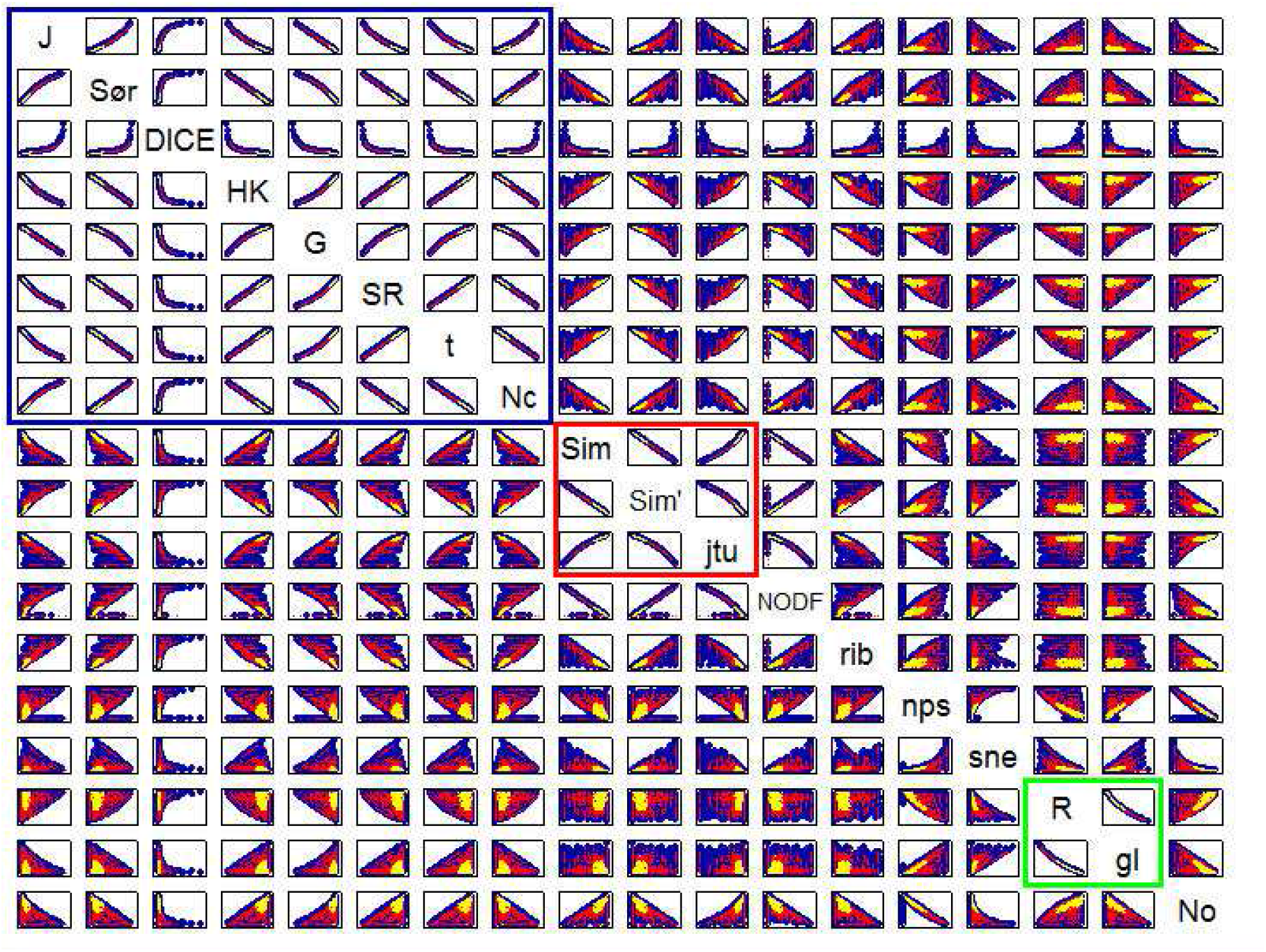
Relationships between the indices for pairs of simulated random (blue dots, N=1352; see Figure R3 in Šizling et al., 2023 for details) and observed (red – plants, N=946; yellow – Ice Shield microbes, N=406) assemblages (Data description R6 in Šizling et al., 2023). The species richness values of assemblages in each pair are uncorrelated, i.e., numbers *S*_*X*∩*Y*_, *b* = *S*_*X*_ − *S*_*X*∩*Y*_ and *c* = *S*_*Y*_ − *S*_*X*∩*Y*_ vary between 0 and 1, they are drawn from a uniform distribution and are mutually s-independent. In *N*_0_and *N*_*NODF*_(i.e., where the equality between *S*_*X*_ and *S*_*Y*_ affects the result) species richness is a random integer between 1 and 20 species. Where possible, the notation is adopted from Gaston et al. (2007). The notation in the plot is simplified. From up to down: *J* – Table 1, *β*_*Sør*_ – Table 1, *β*_*DICE*_ - Raup & Crick (1979), *β*_*HK*_ – Harte & Kinzig (1997), *β*_*G*_ – Table 1, *β*_*SR*_ – Schluter & Ricklefs (1993), *β*_*t*2_ – Table 1, *N*_*C*_ - Wright & Reeves (1992) standardized as in Gotelli & McCabe (2002), *β*_*Sim*_ – Table 1, 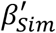 - Table 1, *β*_*itu*_ – Table 1, *N*_*NODF*_– Table 1, *β*_*rib*_ – Ruggiero et al. (1998), *β*_*nps*_ – Table 1 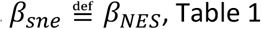, *R* – Table 1; *β*_*gl*_ – Table 1, *N*_0_-Patterson & Atmar (1986) (see Table R1 in Šizling et al., 2023 for index definitions). Blue, red, yellow, and green rectangles delimit the families of Jaccard index, Simpson beta, and richness contrast, respectively.

#### 1. Jaccard index family

The largest family of indices is grouped around the Jaccard index (*J*; Jaccard, 1912). It has 9 indices of (dis)similarity, including Sørensen (*β*_*Sør*_; as defined in Gaston et al., 2007), *β*_*HK*_ (Harte & Kinzig, 1997), *β*_*G*_ (Gaston et al., 2001), *β*_*SR*_ (Schluter & Ricklefs, 1993), *β*_*t*2_ (Wilson & Shmida, 1984) and *β*_*DICE*_ (Raup & Crick, 1979). The Bray-Curtis index of dissimilarity (*β*_*BC*_; Bray & Curtis, 1957), belongs to the Jaccard index family when computed from incidences. A classical index of nestedness, *N*_*c*_ (Wright & Reeves, 1992) also belongs to the Jaccard index family (Figure 5; Thesis T01.20 in Šizling et al., 2023) when standardized by *S*_*X*_ + *S*_*Y*_ (Gotelli & McCabe, 2002). The indices of the Jaccard index family capture the phenomenon of species turnover (Thesis T02 in Šizling et al., 2023). All indices from the Jaccard index family that decrease with increasing *J* (e.g., *β*_*t*2_ or *β*_*Sør*_) are measures of turnover (Figure 5). Whittaker beta-diversity index, when applied to pairwise comparison, also belongs to this family (Tuomisto, 2010b; Table R1 in Šizling et al., 2023).

#### 2. Simpson beta family

The second large family consists of Simpson’s beta (*β*_*Sim*_; Simpson, 1943), Simpson’s nestedness (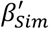 Lennon et al., 2001) and *β*_*itu*_ which was previously referred to as the turnover component (Baselga, 2012). Two classical indices of nestedness, the Discrepancy (*D*; Brualdi & Sanderson, 1999), and the standardized *N*_*C*_ by Wright & Reeves (1992), also belong to the Simpson beta family under certain circumstances. The pairwise index *D* does not belong to any family if we compare just two assemblages. However, if we increase the number of sites being compared, then *D* (standardized as in Greve et al. 2005) belongs to the Simpson beta family, because it converges to the mean value across all *β*_*Sim*_ (Thesis T03 in Šizling et al., 2023). In practice, n of sites >10 guarantees

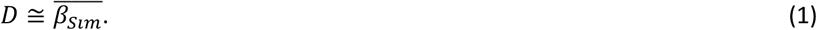

The pairwise *N*_*C*_ also belongs to the Simpson beta family if standardized as suggested by its authors (Wright & Reeves, 1992; Table R1 in Šizling et al., 2023), and if species richness is high (Thesis T04 in Šizling et al., 2023). All the indices from the Simpson beta family capture either the phenomenon of nestedness (Thesis T05 in Šizling et al., 2023), or reversed nestedness (*β*_*Sim*_ and *D*).

#### 3. Richness contrast family

The third family so far consists of just two indices: *β*_*gl*_ (Table 1; Lennon et al., 2001) and *R* (Table 1; Newbold et al., 2016). The *β*_*gl*_ captures the contrast in species richness between two sites (Thesis T06 in Šizling et al., 2023). *R* scales negatively with *β*_*gl*_ and so we call *R* an index of species richness uniformity. Although *β*_*gl*_ is older, the *R* is simpler and thus more tractable than *β*_*gl*_, and so we will use *R* rather than *β*_*gl*_ in our equations(e.g., Equation 2).

#### 4. Other families

There are several other families of indices that do not capture any phenomenon in focus (Thesis T07; Thesis T08; Thesis T09; Thesis T10; Thesis T11; Thesis T12, and Thesis T13 in Šizling et al., 2023). The largest of them consists of *β*_*rps*_ and *β*_*nps*_ (Table 1). Both the indices were introduced by Podani and Schmera (2011) and were called replacement (meaning the one-for-one replacement) and nestedness, respectively. In addition, there are families, each consisting of a single index (Figure 5). These are: Ruggiero index of beta-diversity (*β*_*Rib*_; Ruggiero et al., 1998), Baselga nestedness-resultant components of both Sørensen dissimilarity (*β*_*sne*_; Baselga, 2012; labelled as *β*_*nes*_ in Baselga, 2010) and Jaccard dissimilarity (*β*_*jne*_; Baselga, 2012; not plotted); the classic index of nestedness, *N*_0_(computed for two sites; Patterson & Atmar, 1986), and the *N*_*NODF*_ index of nestedness (computed for two sites; Almeida-Neto et al., 2008). The *N*_*NODF*_ would have belonged to the Simpson beta family if it did not violate the requirement of continuity, which splits the scaling line in the bivariate plot into two different trajectories (Figure 5), which makes any inference difficult.

### i-independent combinations of indices

Although we have delineated nine families of indices, it does not mean that there are nine i-independent indices. The reason is that three indices from any three families are i-dependent even if any single pair from this triplet is i-independent. Only two families are then i-independent with certainty, and the indices of the other families can be calculated from these two (Box 2). A clear case of mutual i-dependence of a triplet of indices is the combination of the Jaccard index family, the Simpson beta family and the richness contrast family. A simple rearrangement of Equation 12 (where *R* = *I* and *k*_1_, *k*_2_, *l*_1_, *l*_3_ = 0, *k*_3_, *l*_2_ = 1, see Box 2), gives

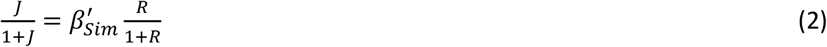

which says that the reversed turnover increases with increasing nestedness and species richness uniformity (reversed contrast). Similarly, for the one-for-one replacement (*β*_*rps*_; Schmera et al., 2020), Equation 12 gives

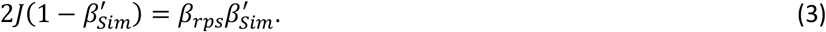

The results (Equation 2, and Equation 3) can be generalized to all the considered indices that can be defined by Equation 8 (Box 2), which includes all dimensionless and continuous indices listed in Table 1 and Table R1 in Šizling et al., 2023, and even indices that have not yet been introduced. This does not include indices that have been produced using partitioning (e.g., *β*_*sne*_), because the partitioning often produces i-dependency (for the algorithm see Box 3, and Thesis T14 in Šizling et al., 2023).

The algorithm in Box 3 provides a tool for converting indices from multiple studies into common terms, assuming that the authors calculated and published at least two i-independent presence-absence indices for their data (e.g., as in Xu et al., 2003).

### Special cases: families of indices when assemblages are interrelated

Species assemblages are often interrelated due to similarity in habitats or due to dispersal. This may lead to two special cases in which the indices that would normally belong to different families appear as i-dependent. First, let us assume an effect that limits variation of species richness between sites. In extreme, *S*_*X*_ = *S*_*Y*_. In this case, there is no species richness contrast, i.e. *R* = 1 and Equation 2 turns into

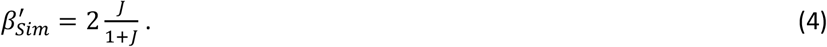

This means that the Jaccard index and the Simpson beta families become apparently i-dependent (Figure R3 in Šizling et al., 2023). Second, assemblages can be perfectly nested, i.e., the species-poor assemblages have no unique species. In this case, *S*_*X*∩*Y*_ = min(*S*_*X*_, *S*_*Y*_) and so 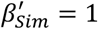 It follows from Equation 2 that

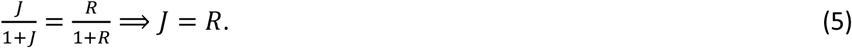

This means that in such special cases the families of the Jaccard index and the richness contrast appear no longer i-independent (Figure R3 in Šizling et al., 2023). Third, pairs of assemblages can have equal proportion of shared species which gives

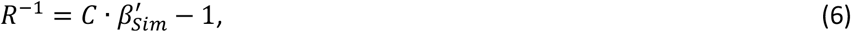

where *C* is a constant. These special cases can easily be recognized in any data.

The apparent loss of i-independences in these special cases is trivial, but it illustrates an important point: Even if indices are i-independent, they appear more i-dependent on each other (i.e., their values begin to cluster along a strictly monotonic relationship and thus become more s-dependent) as species richness values across sites become similar, and/or when assemblages approach complete nestedness (Figure 6).

**FIGURE 6.**
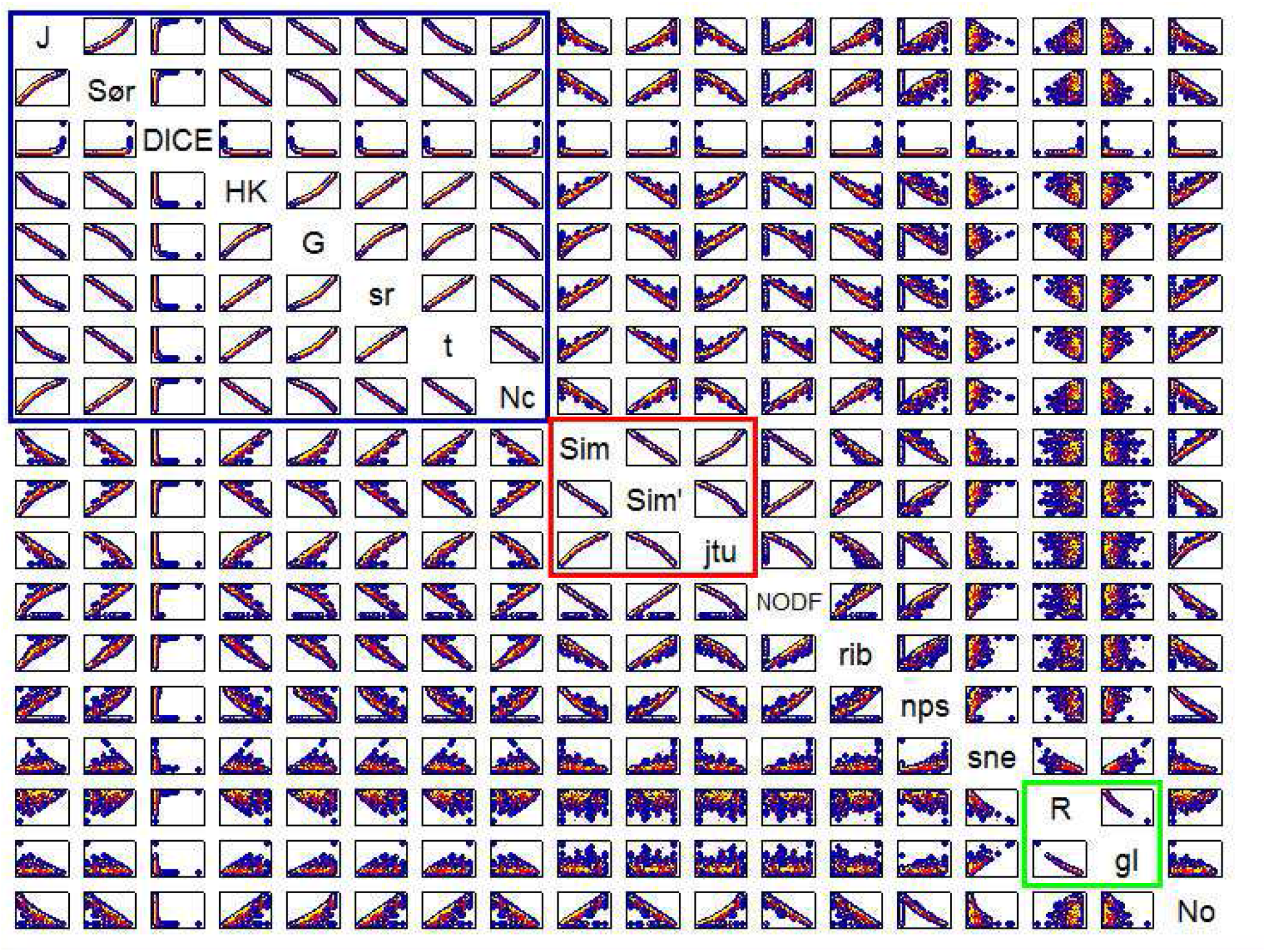
Relationships between pairwise indices of assemblages which have low (0%-20%) species richness contrast (simulations; see Figure 3 in Šizling et al., 2023 for details) or which are spatially close to each other (observed data; Data RI2 in Šizling et al., 2023). Blue (N=1254) and yellow (N=98) dots show data generated by the simulations of random assemblages, and red dots show data for the plant and Ice-Shield microbe assemblages that are closer than 0.5 km from each other (N=98) (for details on data see Data description R6 in Šizling et al., 2023). The cluster of yellow dots is a random subsample of the blue dots, and the N of each subsample (yellow dots) is the same as the N of respective observations (red dots), to allow for comparison. In accord with the theory, when there is low variation in species richness or when the assemblages are close to each other, the Jaccard index, Simpson beta, Ruggiero beta, and *N*_0_families collapse together into one big family of almost i-dependent (and definitely s-dependent) indices. For the meaning of abbreviations see the legend to Figure 5.

### Statistical non-equivalence of the indices

We have argued that the indices within a given family are equivalent, since they are i-dependent and provide the same information. However, they are not equivalent statistically. The reason is that they often scale non-linearly with each other. For instance, within the Jaccard index family there are three groups of indices that scale in a linear manner with each other (group i: *J, β*_*G*_; group ii: *N*_*c*_, *t, β*_*SR*_, *β*_*HK*_, *β*_*BC*_, *β*_*Sør*_; group iii: *β*_*DICE*_; Figure 5), and are statistically equivalent. Between these groups, however, the indices scale nonlinearly (see, e.g., *J* and *β*_*Sør*_ in Figure 5), and this decreases the correlation coefficient between otherwise i-dependent indices, which decreases s-dependence between the indices (e.g., Koleff & Gaston, 2002; Lyashevska & Farnsworth, 2012).

The values of an index within one family that scales in a non-linear manner (Figure 5) with another index can also be seen as transformed values of the other index. For example, the *β*_*t*_ and *β*_*HK*_ represent identical transformations of *J* (Figure 5; Thesis T01.6 in Šizling et al., 2023, and Thesis T01.5 in Šizling et al., 2023). This means that the values of an index and its nonlinear transformation have different frequency distributions, which can affect parametric statistical tests and their sensitivity. This may crucially affect conclusions based on real data with measurement bias (Chao et al., 2005, 2006; Chao & Colwell, 2017).

One option to deal with these issues would be to use an index with the most symmetric/regular frequency distribution in a given system. However, the commonly used indices represent a rather poor spectrum of transformations (Figure 5). We argue that it is better to first pick any index (within a given family) that well describes the phenomenon of interest, and transform/normalize it using an appropriate transformation (e.g., *log*it transformation), rather than pick and choose from the range of existing indices with “good” statistical properties. Alternatively, it is possible to test the index values against a null model, and then the precise distribution of the values is not an issue (Ulrich & Gotelli, 2007, 2013; Chase et al., 2011).

### Consequences of subtracting the indices

Schmera and Podani (2011) showed that only indices with equal denominators are comparable, and thus can be summed or subtracted. The reason is that the denominator scales the nominator and thus it acts as ‘a unit’ in dimensionless indices. However, if we subtract (or sum) the indices with the same denominator, and if these indices are symmetric (sensu Chao et al., 2006) then we necessarily subtract or sum two i-dependent indices, i.e., indices that measure the same phenomenon (see Thesis T15 in Šizling et al., 2023 for details). Since subtracting or summing i-dependent indices leads to another i-dependent index, the operation does not provide any new information. For fundamental reasons why indices of different phenomena cannot be compared and thus summed together nor subtracted from each other see Box 4.

### Inadequacy of partitioning the indices

It makes little sense to develop indices that purify one aspect of diversity patterns, controlling for the other aspects; an example is the attempt to partition “overall beta-diversity” into the nestedness-resultant and turnover components (Baselga, 2010, 2012). The reason is that interdependence between phenomena is an intrinsic characteristic of the phenomena, where different aspects bound each other and removing the interdependence necessarily alters the phenomena itself. For instance, inequality in species richness (species richness contrast) bounds possible values of species turnover.

More specifically, in all situations when we compare two regions or periods, or test for (non)randomness in a diversity pattern with respect to a particular phenomenon, a single index that captures the focal phenomenon is sufficient, regardless of its s-dependence on other indices. We will demonstrate the irrelevance of s-dependence of indices on the triplet 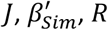. The ideas demonstrated here are then universally applicable, since any pair of i-independent indices is convertible to a pair of indices 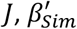, or *R* (Box 3). S-dependence (including relatedness; Chao et al., 2012) between these indices is driven by inequalities:

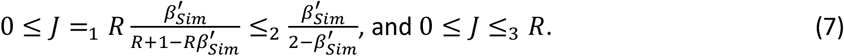

The first equality (=_1_) results from Equation 2, the next inequality ( ≤_2_) involves 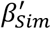 and is a consequence of 0 ≤ *R* ≤ 1, and the last inequality ( ≤_3_) is a consequence of both 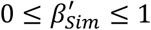 and the first equality (=_1_). The inequality ≤_3_ is well known relatedness between *J* and the contrast between alpha-diversities, which has led to the definition of *β*_*Sim*_ (Table 1). Both bounds ( ≤_2,3_) have been taken as an argument that *J* and *β*_*Sim*_ must be corrected for the effect of the other indices (e.g., Simpson, 1943; Baselga, 2010). We argue that, when we are interested in turnover (not nestedness nor other phenomena), then the indices of the Jaccard family must not be corrected for variation in species richness contrast, regardless of any interdependence between these phenomena. The rationale is that (i) the *J* is a measure of reversed turnover regardless of its relatedness with R (i.e., *J* meets the reversed constraints of turnover in Figure 3; Thesis T15 in Šizling et al., 2023), and that (ii) the bounds ( ≤_2,3_) only show the limits of turnover given the level of nestedness or uniformity (*R*) in species richness across sites. Obviously, it would be possible to use the value of *J* relative to its maximum (≤_1_), but this value reflects just the level of uniformity *R* (Figure 7), so using this relative *J* is equivalent to direct measurement of *R*.

**FIGURE 7.**
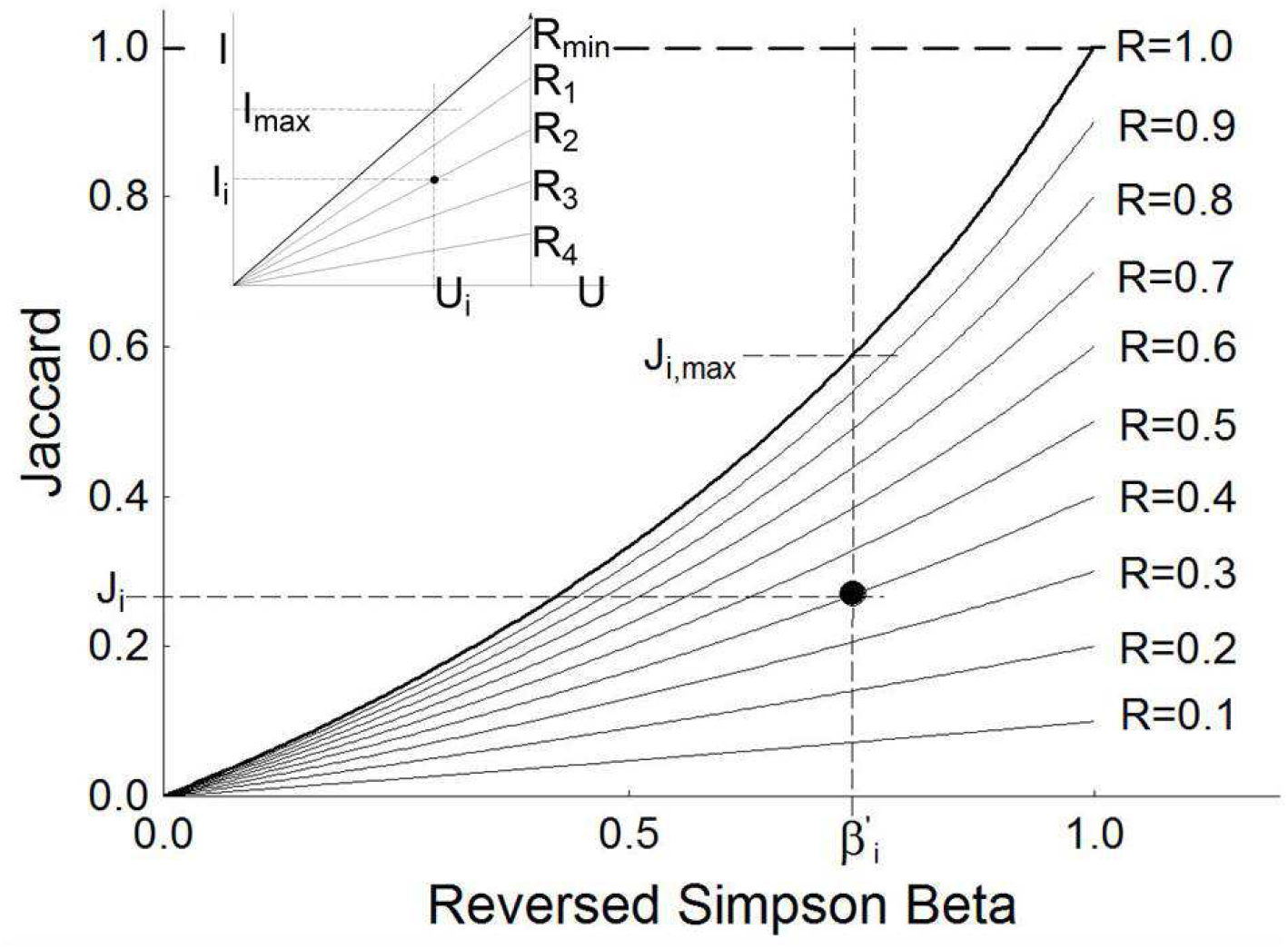
The reason why mutual dependence of indices cannot be eliminated to purify individual effects. The relationship between Jaccard similarity ( *J*) and Simpson nestedness 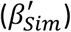 is determined by species richness uniformity R (Equation 7), each line shows points with equal R (the dot has R=0.4). *J* is limited by 1 (dashed bold line) and each particular *J*_*i*_ is limited by *J*_*i*,*max*_, which is a function of 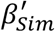 (bold line, Equation 7). The value of *J*_*i*_ relative to the limit of one should be read as similarity (or reversed turnover) without a need for any partitioning of *J* or 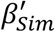 into components. We say that *J* is a similarity *per se* (or reversed turnover *per se*). The value *J*_*i*_ rescaled by *J*_*i*,*max*_ shows *R*_*i*_, and should be read as uniformity *per se* (or reversed species richness contrast *per se*). The fact that *R* affects *J* is a matter of i-dependence between triplet 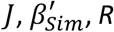, and thus the interplay between *J* and *R* should not invalidate their unique meaning determined by the constraints (section *Formal definitions of the phenomena based on their constraints*). This is similar to the Ohm’s law in physics where the triplet of voltage (*U*), current (*I*), and resistance (*R*) are mutually dependent (*U=RI*; see inset). The understanding in physics is that *I*_*i*_ is the current regardless of its dependence on *R* and *U*. No one would relativize *I* to *R*, nor say that in the absence of voltage *U* equals *I*, and thus *U* minus *I* measures the ‘voltage resultant component’. In the same way, *J*_*i*_ is reversed turnover *per se* even though it also scales with reversed richness contrast *per se*. For the same reason 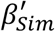 is a measure of nestedness *per se*, without a need for partitioning into components of turnover and/or nestedness.

A general intuition illuminating the inadequacy of the partitioning of indices (defined by Equation 8) stems from the realization that any arrangement of assemblages is fully described by two i-independent indices (plus species richness as a scaling parameter). This implies that if we have two i-independent indices, there are no degrees of freedom for residual information about the assemblage arrangement which would follow from the partitioning. A partitioned index that combines two i-independent indices thus does not measure a phenomenon purified from the effect of the other phenomenon, but a combination of the phenomena, which is a different phenomenon. A good example is the attempt of Simpson (1943) to purify turnover from species richness contrast, which has led to 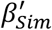, i.e. the index which directly measures nestedness (Almeida-Neto et al., 2008). Any partitioning thus leads either (1) to an index that measures one of the phenomena we already measure by another established index (as in the above case) or (2) an index that is not strictly monotonically related to any of the phenomena that we want to measure (as in Baselga′s approach in Figure 1).

### Empirical examples: How to make inference from the indices?

To provide real world examples of an inference based on similarity indices, we use data from Šizling et al. (2016) who studied temporal changes of Central European plants during Holocene, and Xu et al. (2023) who reported global patterns of angiosperm trees. The main purpose of these examples is to show that the increasingly popular partitioning of these indices (Baselga, 2010, 2012) may lead to interpretations that do not correspond to any of the four phenomena intuitively distinguished by ecologists (section *Predefinitions of four diversity phenomena*).

#### Example 1: Temporal change in Holocene plants

Šizling et al. (2016) reported a temporal increase in mean species richness ⟨*S*⟩ from 27 to 32, almost no change of 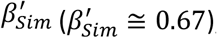, and a steep increase of average *J* from ca. 0.4 to ca. 0.5 during the short period (5,900-5,700 BP) when agriculture was introduced in the region. *J* and 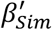 uniquely determine *R* (Equation 2), which increased from 0.74 to 0.99 (Figure 8). In addition, *R* uniquely determines the contribution of species-poor (*S*_*poor*_ = ⟨*S*⟩ 2*R*⁄(*R* + 1)) and species-rich (*SR*_*rich*_ = *R*^−*l*^. *S*_*poor*_) sites to the average alpha diversity (Thesis T16 in Šizling et al., 2023). *S*_*poor*_ increased from ca. 23 to 32, and *SR*_*rich*_ increased from ca. 31 to 32. We conclude that recent plant assemblages in Central Europe show lower spatial turnover (Figure 8c), and higher uniformity (Figure 8c) than those before the introduction of agriculture. Since the average nestedness (Figure 8c) hasn’t changed from before agriculture until today, the species poor sites have also kept a constant ratio of unique to common species. The introduction of agriculture was therefore accompanied by homogenization of plant assemblages, with proportionally higher increase of richness at species-poor sites, and increasing average species richness of all sites. This homogenization of assemblages as a consequence of agriculture is in agreement with other analyses (e.g., Kolář et. al., 2022) and corresponds to the visual inspection of Figure 8a.

**FIGURE 8.**
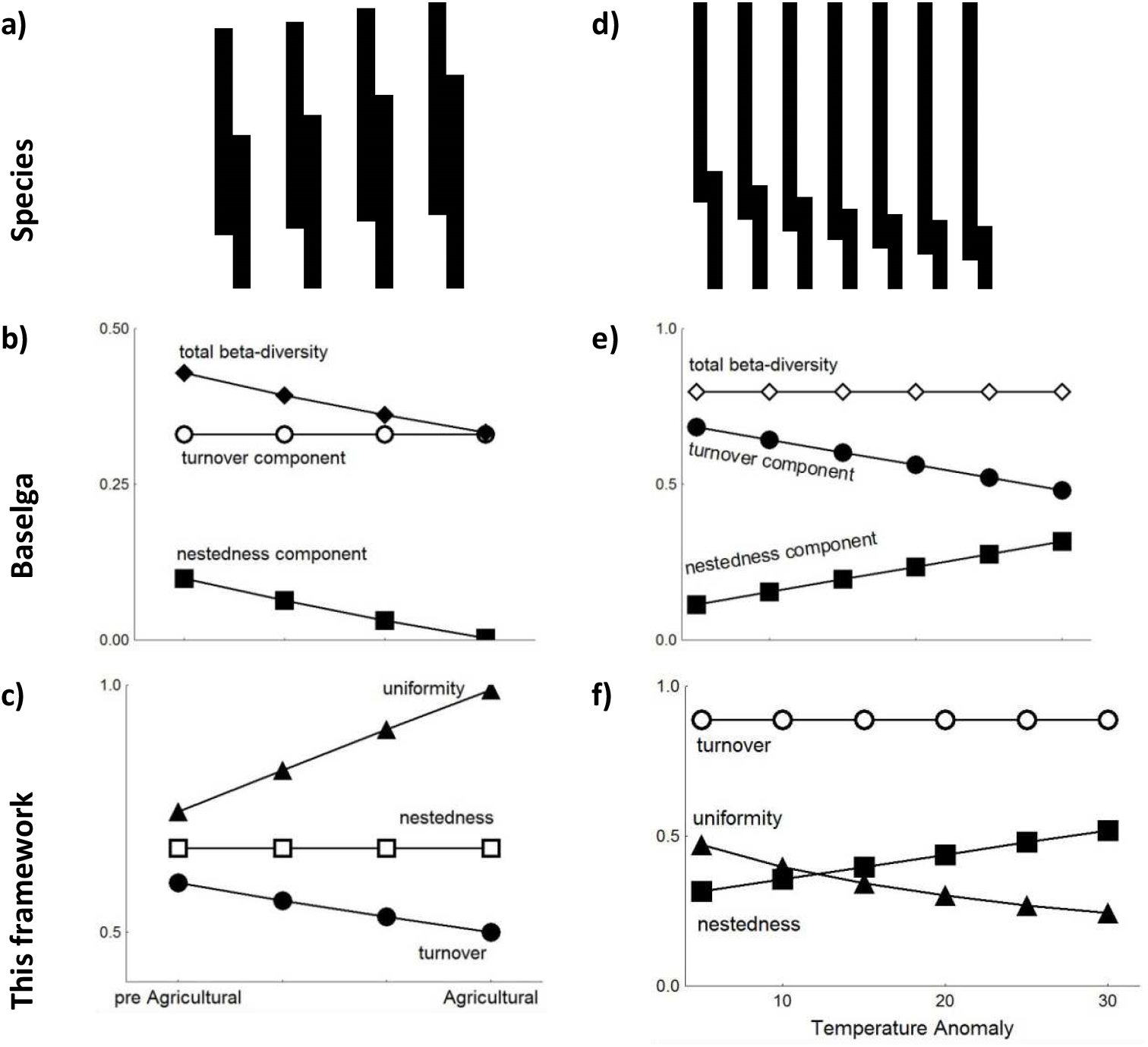
The illustration of the difference between the approach that uses Baselga’s (2010) additive partitioning of pairwise indices (b,e) and the framework presented in this paper (c,f), using empirical data from Šizling et al. (2016, left) and Xu et al. (2023, right). Panels (a) and (d) show the species assemblages and respective overlaps in the lists of species (see caption to Figure 1). We used the algorithm in Box 2 and Box 3 to compute (a) and (b) from (c), and (f) and from (e), as only data in (c) and (e) was published (the computed *S*_*X*_ ranges from 75.62 to 89.54; *S*_*Y*_ ≅ 35.66 to 21.74 and *S*_*X*∩*Y*_ ≅ 11.28 to 11.28 in Xu. et al. 2023, *S*_*X*_ ranges from 30.97 to 32.16; *S*_*Y*_ ≅ 23.03 to 31.84 and *S*_*X*∩*Y*_ ≅ 15.43 to 21.33 in Šizling et al. 2016). The figures (b) and (e) display trends for *β*_*Sim*_ (circles), *β*_*sne*_ (squares), and *β*_*t*2_ (diamonds), labeled according to Xu et al. (2023) as turnover component, nestedness component, and total beta-diversity, respectively (*β*_*Sim*_ + *β*_*sne*_ = *β*_*t*2_). The figures (c) and (f) show trends for 1 − *J* (circles), 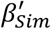 (squares), and *R* (triangles), labeled according to this report as turnover, nestedness, and uniformity, respectively. Differences between (b) and (c), as well as (e) and (f), lead to divergent interpretations when applying our framework compared to Baselga’s (2010, 2012). The lines show least square regressions; where the regression was not significant (*α* > 0.05; open symbols) we plotted a constant line.

When we apply Baselga’s (2012) partitioning method to the data of Šizling et al. (2016), we get *β*_*sne*_ = 0.1, *β*_*Sim*_ = 0.33 for the pre-agricultural landscape and *β*_*sne*_ = 0.00, *β*_*Sim*_ = 0.33 for early agricultural landscape. This reads in the Baselga’s (2010, 2012) framework as no temporal change of “turnover component”, and decreases in both the “nestedness-resultant component” and total beta-diversity (Figure 8b). This interpretation based on Baselga’s (2010, 2012) partitioning thus directly contradicts our framework (Figure 8c). Baselga’s framework misses the increasing uniformity of species richness (due to the increase of species richness of species-poor assemblages) which seems to be driving the other observed patterns.

#### Example 2: Global patterns of trees

Xu et al. (2023) used the Baselga (2010) framework and reported an increase in the regression line of the nestedness-resultant component (*β*_*sne*_) with an increasing temperature anomaly from ca. 0.11 to 0.32, and a decrease in the regression line of the turnover component (*β*_*Sim*_) from ca. 0.68 to 0.48 (Figure 8e). The total beta-diversity (*β*_*sne*_ + *β*_*Sim*_) did not show a significant trend along the temperature anomaly gradient.

To reanalyze Xu et al.’s (2023) data, we used our framework (Box 2; Box 3, and Calculator RI4 in Šizling et al., 2023) and computed the average *S*_*X*_, *S*_*Y*_, and *S*_*X*∩*Y*_ that correspond to the indices reported by Xu et al. (2023; Figure 8e). Since Xu et al. (2023) did not report richness, we standardized it to *S*_*X*_ + *S*_*Y*_ − *S*_*X*∩*Y*_ ≅ 100 without loss of generality. Our reanalysis of the data from Xu et al. (2023) suggests no significant trend in turnover (in contrast to a significant decrease from Baselga’s framework), increasing nestedness (in line with Baselga’s framework), and decreasing richness uniformity with an increasing temperature anomaly (Figure 8f). This indicates a heterogenization of assemblages as the temperature anomaly increases. Visual inspection of the arrangement of species assemblages (Figure 8d) indicates the pattern of increasing contrast in species richness of both assemblages (i.e. decreasing uniformity) associated with increasing nestedness, and no pattern of species turnover (as the overlap is constant). This agrees with our interpretation of the indices, but is in striking contrast to the interpretation within Baselga’s framework.

### Practical guidelines

Practical inference from pairwise indices can have different purposes (Anderson et al., 2010), for instance: (i) exploration of diversity (e.g., Qian et al., 2009) or (dis)similarity (e.g., Simpson, 1943) between biotas of two sites or regions, (ii) revealing non-random origin of a spatial or temporal biodiversity patterns (e.g., Patterson & Atmar, 1986; Ulrich & Gotelli, 2013), (iii) meta-analysis based on indices extracted from the literature, and (iv) exploration of the behavior of the indices along temporal, spatial or environmental gradients. Based on previous considerations, here are practical recommendations for using the presence-absence indices:

1. Avoid partitioning of indices that supposedly removes an effect of one phenomenon from an effect of another phenomenon (e.g., purifying turnover from the effect of richness contrast or purifying nestedness from the effect of turnover). These indices are either mathematically flawed (e.g., they violate the requirement of continuity), and their meaning is thus unclear, or they measure a different phenomenon from what they were originally claimed to measure. An example of the latter is the standardization of an index relative to its maximum value. For instance, standardization of Jaccard index by its maximum determined by the nestedness (bold line in Figure 7) is in fact a measure of species richness contrast (Figure 7). The reason is that nestedness, together with richness contrast *R*, uniquely determine *J*, so that measuring *J* relative to *Jmax* (bounded by nestedness) is equivalent to measuring *R*.
2. When choosing an index, first consider which phenomenon it should capture (Table 1; Table 2), then select a corresponding family of indices (see Figure 5; Table R1 in Šizling et al., 2023; Manual RI3 in Šizling et al., 2023, and Calculator RI4 in Šizling et al., 2023). It does not matter which index within the family is selected, as all indices within any family can be converted to each other and are thus practically equivalent. If there is a need for an appropriate statistical distribution of the index, a proper transformation is preferable to an invention of, or search for, a new index.
3. When comparing already published indices, use the equations in Table R1 in Šizling et al. (2023) or the Calculator RI4 in Šizling et al. (2023) to convert them to a common index that is the most desirable for a given purpose. If a desirable common index is not available in the publication, the desirable index can be calculated from any i-dependent index, or any two i-independent indices (Box 2 and Box 3). The indices of two different phenomena (two i-independent indices) are never commensurable.
4. Use null models that randomize species incidences to assess the statistical significance of the phenomenon in focus (Gotelli & Ulrich, 2012). Null models are useful for distinguishing inherent correlations (Chao et al., 2012) from ecologically meaningful correlations and relationships. Note, however, that relevant null models are still under development (e.g. Chase et al., 2011; Legendre, 2019), and it may not always be clear what exactly to randomize in the site-by-species matrix.
5. When publishing indices from your research, ensure that these indices conserve complete information, i.e., publish at least two i-independent indices (*J*, and *β*_*Sim*_ or *R* are recommended) plus regional and mean local species richness, if available. Use our Calculator RI4 in Šizling et al. (2023) when in doubt about whether the employed indices carry complete information.

## Discussion

We demonstrated that the phenomena that are measured by different indices constrain each other at the conceptual level, and respective indices are thus necessary related sensu Chao et al. (2012). This leads to their statistical dependence. It is pointless to search for s-independent indices, because any effort to eliminate s-dependence (or relativize the index in focus) results in a measure of a different phenomenon. In other words, there are no s-independent indices that capture different, but necessarily mutually dependent phenomena.

**TABLE 2.** Table of possible inference from the variation of pairwise indices along temporal, geographic, or environmental gradients. Some inference can be based on variation of a single index, regardless of other indices (first three rows). However, more specific inference follows from situations when one index changes along the gradient, while the other remains constant or shows only negligible variation (all remaining rows).

| Variation along a gradient |  | Inference |
| --- | --- | --- |
| Yes | No |  |
| $J$ | | Change in turnover between regions/periods, change in proportion of shared species relative to species richness of both assemblages |
| $\beta'_{Sim}$ | | Change in nestedness compared between regions/periods (i.e., nestedness for region 1 vs nestedness for region 2), change in proportion of common species relative to richness of species poorer sites. |
| $R$ | | Change in species richness contrast (uniformity) between regions/periods; change in variation of alpha diversity |
| $\beta'_{Sim}$ | $S_{rich}$ | Change in nestedness between regions/periods is driven by exchange of species at species poor sites |
| $\langle S \rangle$ | $S_{rich}$ | Difference of alpha diversity between regions/periods is driven by sites with fewer species |
| $\langle S \rangle$ | $S_{poor}$ | Difference of alpha diversity between regions/periods is driven by species-richer sites |
| $R$ | $\langle S \rangle$ | Regions/periods differ in spatial variation of alpha diversity without change of its average value; |
| $\beta'_{Sim}$ | $J$ | change in nestedness but not turnover; simultaneous change in lists of species that are unique to species-richer and species-poorer sites; variation in $\beta'_{Sim}$, no change in $J$, and Equation 2 induce necessarily a change in $R$ |
| $J$ | $\beta'_{Sim}, S_{poor}$ | Change in turnover but not nestedness; change in list of species that are unique to species-richer sites; necessarily a change in $R$ |

On the other hand, i-independent indices are desirable, as they provide (by definition) different information about the system. However, a sufficient number of i-independent indices have already been invented (and any not yet invented index is i-dependent on *J* and *β*_*Sim*_, or *R*; Box 2 and Box 3), and there is thus no need for new indices. We have mathematically proven that two i-independent indices, in combination with a species richness value (which acts as a scaling parameter), provide complete information about the assemblages. This is not true for the partitioned indices because the partitioning often reduces the amount of information. Moreover, when variation of species richness across sites is negligible, or when there are no species that are unique to the species-poorer assemblage (i.e., perfect nestedness), i-independent indices appear as if they were i-dependent, and can be derived from each other. The interrelation between assemblages which leads to these effects thus leads to higher correlations between the observed index values than what would follow from the theory (Ulrich et al., 2017), which makes some families of indices almost indistinguishable (Figure 6; Figure R3 in Šizling et al., 2023). This is apparently the reason why it has been so difficult to achieve agreement regarding which indices characterize different phenomena (Lennon et al., 2001; Gaston et al., 2007; Ulrich & Gotelli, 2007; Tuomisto, 2010a; Tuomisto, 2010b; Baselga, 2012; Podani & Schmera, 2011; Ulrich et al., 2017; Schmera et al., 2020).

In terms of practical utility of the indices, the finding that the phenomena distinguished by ecologists are mutually dependent and that two i-independent indices fully characterize the system implies that it is reasonable to calculate just two indices belonging to different families (see section *Results*) to make a proper inference. These indices (for example *J* and *R*) can be, if necessary, converted to any other index within their respective family (and vice versa). However, we have shown that indices from different families cannot be directly compared to each other, as their values characterize different phenomena. Different indices within a family of i-dependent indices are different transformations of each other and their particular values thus reflect just different rescaling.

We have focused only on pairwise indices that compare spatially or temporarily non-overlapping assemblages and have avoided the topic of beta-diversity *sensu stricto* (Whittaker, 1960), which comprises the relationship between local and regional species richness, or, more specifically, between alpha and gamma diversity. This relationship is to some extent related to turnover, since when alpha is considerably lower than gamma, there must be high turnover among communities. However, the precise mathematical links between pairwise community turnover and Whittaker’s beta is a separate issue (Koleff et al., 2003; Tuomisto, 2010b; Chao et al., 2012) and there is no straightforward way to derive turnover from beta-diversity *sensu stricto* for more than two assemblages (Šizling et al., 2011).

It is fundamental that two i-dependent indices cannot quantify two different phenomena. An index that is i-dependent on another index can be viewed as its rescaled version, and no rescaling can affect the phenomenon, since rescaling can be done at any time after the observation. This is in contrast to Tuomisto (2012) who defines ‘conceptual independence’ of indices (Box 1) to delimit her phenomena. In our framework, the phenomena are defined more broadly using the constraints in Figure 3. If two indices obey the particular set of constraints, they quantify the same phenomenon. Mutually i-dependent indices necessarily satisfy the same constraints and thus they quantify the same phenomenon.

Our findings regarding the pairwise comparison of two assemblages can also be applied to pairwise comparisons of two species ranges, and generally to all pairwise indices of interspecific spatial association derived from binary data (Keil et al. 2021). In particular, *S*_*X*_, *S*_*Y*_, and *S*_*X*∩*Y*_ can represent the number of sites occupied by the first species, second species, and both species together, respectively. For instance, the co-occurrence of two species over multiple sites (Diamond, 1975; Connor & Simberloff, 1979; Diamond & Gilpin, 1982; Gotelli & McCabe, 2002) can be measured by the Jaccard index (Thesis T17 in Šizling et al., 2023). Analogically, indices from the Simpson beta family and richness contrast family can be used for characterizing nestedness of species’ geographic ranges (Šizling et al., 2009) and the contrast in species extent of occurrence.

The relationships between indices and families have important implications for understanding the forces shaping distance decay (a strict increase or decrease of an index with increasing distance) in assemblage similarity (Nekola & White, 1999). A direct consequence of Equation 2 is that distance decays of nestedness 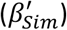 and reversed turnover (measured by *J*⁄(1 + *J*)) follow the same mathematical function when there is no distance decay in the species richness contrast. This happens only in environments without geographical gradients that affect the species richness, for example, the gradient in productivity (e.g., Willig et al., 2003; Currie et al., 2004; Bohdalková et al., 2021). If there are richness gradients, then spatial structuring of the assemblages is anisotropic, and there are different rates of distance decay in different directions. Even more importantly, it is meaningless to assume a universal functional form of the distance decay of community similarity, e.g., exponential (Nekola & McGill, 2014). This follows from the mutual non-linear scaling between different indices, even between the i-dependent ones. Consequently, if one index reveals, say, exponential distance decay, another index can reveal a non-exponential distance decay.

Our results are relevant for the interpretation of hundreds of empirical assessments that used the subtractive partitioning of pairwise indices. To date, the original publication describing the partitioning to nestedness and turnover components (Baselga, 2010) has 3,365 citations, the publication announcing the R package betapart which does the partitioning (Baselga & Orme, 2012) has 1,884 citations, and the R package on CRAN (Baselga et al., 2018) has 852 citations (Google Scholar accessed on 3^rd^ of April 2024). Most of these are empirical studies, including some high profile ones in top journals (e.g., Molinos et al., 2016; Gotelli et al., 2017; Rocha et al., 2018; Blowes et al., 2019; Chase et al., 2020; Xu et al., 2023). The approach has also been gaining momentum in young fields, for example in microbial ecology (Shade et al., 2013). Based on the conceptual problems that we have described, we call for a critical re-evaluation of the practice. A need may also arise for re-analyzing and re-interpreting some of the studies; in this effort, our equations from Box 2, and Box 3 (or Calculator RI4 in Šizling et al., 2023) can be helpful.

In conclusion, our framework systematically deals with problems that unnecessarily generated new indices, and it resolves old issues concerning the mutual dependence of the indices. Based on the distinction between two types of dependence (i- vs s-dependence), we offer a tool for making inference using classical indices, a tool that can be further developed when new spatial or temporal phenomena are identified.

## Acknowledgements

Special thanks to A. Bryson and G. Ridder for help with English, F. He and W. E. Kunin for their comments on the first versions, Z. Sokolí čková, E. Šizlingová for her help in the field, J. Klimešová and A. Bernard for lessons in Arctic plants, and R.K. Colwell and anonymous reviewers for insightful comments. The project was supported by the Czech Science Foundation (EXPRO 20-29554X and GA ČR 19-21341S). We used the Czech Scientific Infrastructure of the University of South Bohemia, J. Svoboda house on Svalbard – CzechPolar2 LM2015078 supported by MŠMT. PK was funded by the European Union (ERC, BEAST, 101044740). Views and opinions expressed are however those of the author(s) only and do not necessarily reflect those of the European Union or the European Research Council Executive Agency. Neither the European Union nor the granting authority can be held responsible for them.

## Author Contributions

AS, DS, and ET conceived the ideas. AS and DS worked out the definitions, concepts, and proofs. ET, JDŽ and ALŠ collected and provided the empirical data. ALŠ did the analyses, simulations, and coded the software tool. ALŠ, DS and PK led the writing with contributions from ET and KMCT.

## BOX 1

**Key concept: The informational dependence of indices**

Information content of two indices is identical when each value of the first index uniquely determines the value of the second index, and each value of the second index uniquely determines the value of the first index (Orlitsky, 2003). We thus define informational dependence (i-dependence) of two variables as a situation when these variables are strictly monotonic transformations of each other (Figure 2). A strict monotony of two variables implies equality of their informational values, and when two indices have equal information content, they cannot capture different phenomena.

In contrast, the kind of independence that is used in ecological literature is almost always of statistical nature, and we call it *s-dependence*. While i-independence refers to a mathematical formula, s-independence refers to data values (i.e., a conditional probability of measuring a particular value of one variable, given a value of another variable).

Chao et al. (2012) distinguished statistical dependence from a dependence of mathematical formulas of indices; two indices are statistically dependent when one index value affects probability density of values of another index. Chao et al. (2012) use the term *relatedness* for dependence of the formulas, and defined it as a situation where the minimum or maximum of one index is affected by the variation of another index. Relatedness, however, is one of the mechanisms that induces s-dependence, since a change in the minimum and maximum possible value of an index has an effect on its probability density, which is the definition of statistical dependence (Figure 2). Importantly, relatedness says nothing about equal or different information carried by the variables in focus. For example, *J* and 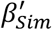 are i-independent, while their values are s-dependent (their values are correlated) because their definitions are related sensu Chao et al. (2012) (Figure 4a,b).

Tuomisto (2012) defines the independence of indices based on different defining formulas. According to Tuomisto (2012), two different formulas, e.g. *I*_1_ = (*A* − *B*)/*B* and *I*_2_ = *A*/*B*, are conceptually independent of each other and quantify two different phenomena. The reason is that *I*_1_ reads as ‘relative difference’ but *I*_2_ reads as ‘proportion’. However, in our termino*log*y *I*_1_ and *I*_2_ are i-dependent and define different measures of the same phenomenon, because *I*_1_ = *I*_2_ − 1, and whatever can be inferred from *I*_1_ can also be inferred from *I*_2_.

## BOX 2

*How many indices do we need?*

For a given system, a multitude of indices can be calculated. What is the minimum number of indices that characterize a given system fully? Here we show that three i-independent indices (including species richness) are sufficient for full characterization of the system, so that any other index can be reconstructed using this information. We adopt the term of independence from mathematics, which is independence of equations in terms of their solutions (the i-independence from Box 1). For this purpose, we need to see definitions of the indices as equations to solve. This allows for two statements: (i) all the focal indices listed in Table R1 in Šizling et al. (2023) are i-dependent on the pair of Jaccard index, *J* (Table 1) and Simpson nestedness, 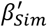 (Table 1); and (ii) all information concerning the difference between two assemblages is captured by 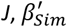, and an index that is i-dependent on species richness (e.g., species richness itself). Here we show why these statements hold.

All the focal indices can be defined as a ratio of two linear functions (see also Table R1 in Šizling et al., 2023):

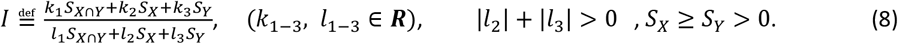

For *J* and 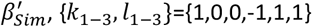 and {1,0,0,0,0,1}, respectively. Following the *log*ic: 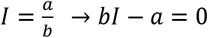, Equation 8 can be converted into the linear equation

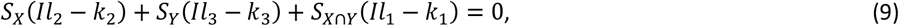

where *S*_*X*_, *S*_*Y*_, *S*_*X*∩*Y*_ are the unknowns and the *I* is a particular value of an index. Equation 9 turns into

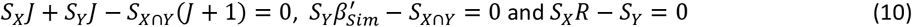

for 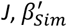 and *R*, respectively. The two equations for two indices (Equation 10), and the three unknown variables (*S*_*X*_, *S*_*Y*_, *S*_*X*∩*Y*_), do not provide a unique solution, and have zero at the right side. If the third equation had zero on the right side, the system would either provide multiple solutions or the only solution would be zero (*S*_*X*_, *S*_*Y*_, *S*_*X*∩*Y*_ = 0, 0, 0). We therefore need an independent equation with a nonzero right side to get unique information on *S*_*X*_, *S*_*Y*_, *S*_*X*∩*Y*_. This equation is

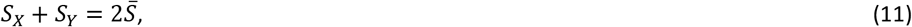

where 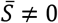 is the expected alpha diversity (for the proof that Equation 11 is i-independent of Equation 10; see Thesis T18 in Šizling et al., 2023). The three equations (any two equations from Equation 10, and Equation 11) determine *S*_*X*_, *S*_*Y*_, *S*_*X*∩*Y*_ uniquely, and thus no other index (even if its definition does not follow Equation 8) can further specify the solution. Moreover, if we focus on the indices which can be expressed using Equation 8, we can compute the value of any other index from the values of *J* and 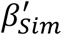 or *R* as

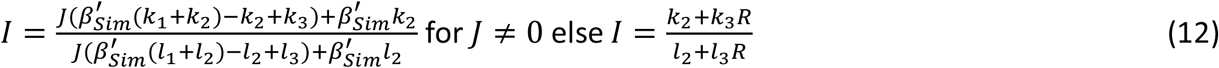

where *k*_*i*_, *l*_*i*_ define the new index *I* (for the proof see Thesis T19 in Šizling et al., 2023).

## BOX 3

*How to unify indices from across the literature*

A large number of different indices are spread over the literature (e.g., Gaston et al., 2007; Baselga, 2012; Schmera et al., 2020). However, if one wishes to do a meta-analysis and compare indices from different published sources, one needs to make the indices comparable, and to convert them to only one or two reference indices. Here we show how this can be done, based on the idea that an index is defined by an equation. Firstly, we need a publication that utilizes at least two i-independent indices, *I*_1_, *I*_2_, that are expressed as in Equation 8 (see Box 2). Then we write a system of three independent equations: two equations Equation 9, each for one of the indices, and one scaling equation Equation 11. If 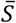 is missing from the publication, we can put 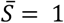 without loss of generality. The reason is that the indices to be converted (Equation 8) are i-independent of species richness and therefore the exact value of 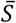 does not matter in the case. The third equation then is

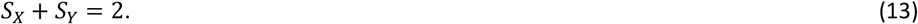

The solution of the three equations (see *Gauss Elimination Method*) is *S*_*Xcomp*_, *S*_*Ycomp*_ and *S*_*X*∩*Ycomp*_. This solution is different from the solution based on the original data behind the published source, but we can get the original average values *S*_*Xorig*_, *S*_*Yorig*_, *S*_*X*∩*Yorig*_ by simple rescaling of the computed values (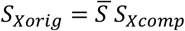 and so on), in case we know ⟨*S*⟩. From *S*_*Xcomp*_, *S*_*Ycomp*_, *S*_*X*∩*Ycomp*_ we can compute any index that is i-independent of species richness, even for an index that cannot be expressed as Equation 8 (e.g.,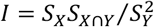). Some special cases are the indices that originated from additive partitioning (*I*_1_ = *I*_*a*_ − *I*_*b*_) such as *β*_*sne*_, *β*_*jne*_, of *β*_*nps*_ (Table 1). These indices cannot be universally expressed as Equation 8, and their equations are no longer linear. In the case of additive partitioning, we can compute *I*_*b*_ from two i-independent indices *I*_1_ and *I*_2_ using Equation 14

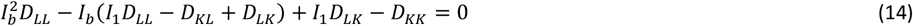

where *D*_*XY*_ are determinants listed in Thesis T14 in Šizling et al. (2023). *I*_*a*_ then follows *I*_1_ + *I*_*b*_. If *I*_*a*_, *I*_2_, or *I*_*b*_, *I*_2_, or *I*_*a*_, *I*_*b*_ are i-independent then we can follow the above algorithm that uses linear equations and compute any even not yet invented index. However, as Equation 14 may have two realistic solutions (both solutions are within minimum and maximum possible range of *I*_*b*_), the partitioning often leads to the loss of information. For example, if Šizling et al. (2016) had only reported two partitioned indices (e.g., *β*_*sne*_ = 0.1 and *β*_*nps*_ = 0.61 for pre-agricultural landscape, and *β*_*sne*_ = 0.0 and *β*_*nps*_ = 0.51 for agricultural landscape, it would not be sufficient to infer the homogenization of assemblages. This is because these two indices do not distinguish between an increase or decrease in uniformity (R), as R could have changed from 0.74 to either 1 or 0.32, respectively (see Thesis T20 in Šizling et al., 2023 for further details).

## BOX 4

*Different phenomena are incomparable*

There are several reasons why indices of different phenomena are incomparable. One of them is that the relationship between two phenomena (e.g., turnover and nestedness) is modulated by an index of a third phenomenon (e.g., species richness contrast), which is necessarily i-dependent on the two phenomena in comparison. This interdependence means that the indices cannot be viewed in isolation (Table 2). An example of the relationship of three indices is Equation 2, which can be transformed to involve any triplet of similarity indices (see Box 2 for the algorithm). We cannot put the equation sign nor inequality sign between the indices of nestedness and turnover without taking into account the effect of species richness contrast (Podani & Schmera, 2012). It is not 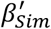 but the product of 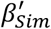 and *R*/(*R* + 1) that is comparable with *J*/(*J* + 1). Similarly, no one would directly compare *U* with *I* in Ohm’s law, because the relationship between *U* and *I* is modulated by *R, U* = *RI* (Figure 7).

The most fundamental reason, however, stems from the theory of measurement (ISO, 2009; Mari et al., 2023), which states that: (i) there are different concepts of measurement (kinds of quantities) that are incomparable even though they may have the same units (e.g., torque and work are incomparable but have the same unit, the Newton meter), and that (ii) the dimensionless quantities are not commensurable if they capture different phenomena.

The phenomena addressed here capture different concepts of similarity between assemblages (they are different kinds of quantities) and are therefore incomparable. The reason is that they are defined using different sets of constraints (Figure 3). In particular, turnover is conceptually defined as species overlap (*S*_*X*∩*Y*_) relative to the richness of both assemblages (*S*_*X*_, *S*_*Y*_) (e.g., relative to total richness in the case of Jaccard, and mean richness in the case of Sørensen index). Unlike turnover, nestedness is conceptually defined only by the property of the poorer assemblage (min(*S*_*X*_, *S*_*Y*_), *S*_*X*∩*Y*_), which makes it i-independent of the species richness of the richer assemblage (max(*S*_*X*_, *S*_*Y*_))). This means that the index of turnover must include both *S*_*X*_ and *S*_*Y*_ in the denominator (Thesis T21 in Šizling et al., 2023), whereas the denominator of the index of nestedness can include either *S*_*X*_ or *S*_*Y*_, but not both. Different denominators make the indices of turnover and nestedness incomparable. This incomparability of different phenomena implies that their quantities cannot be summed nor subtracted.

## RI1: Repository Information

**TABLE R1.**
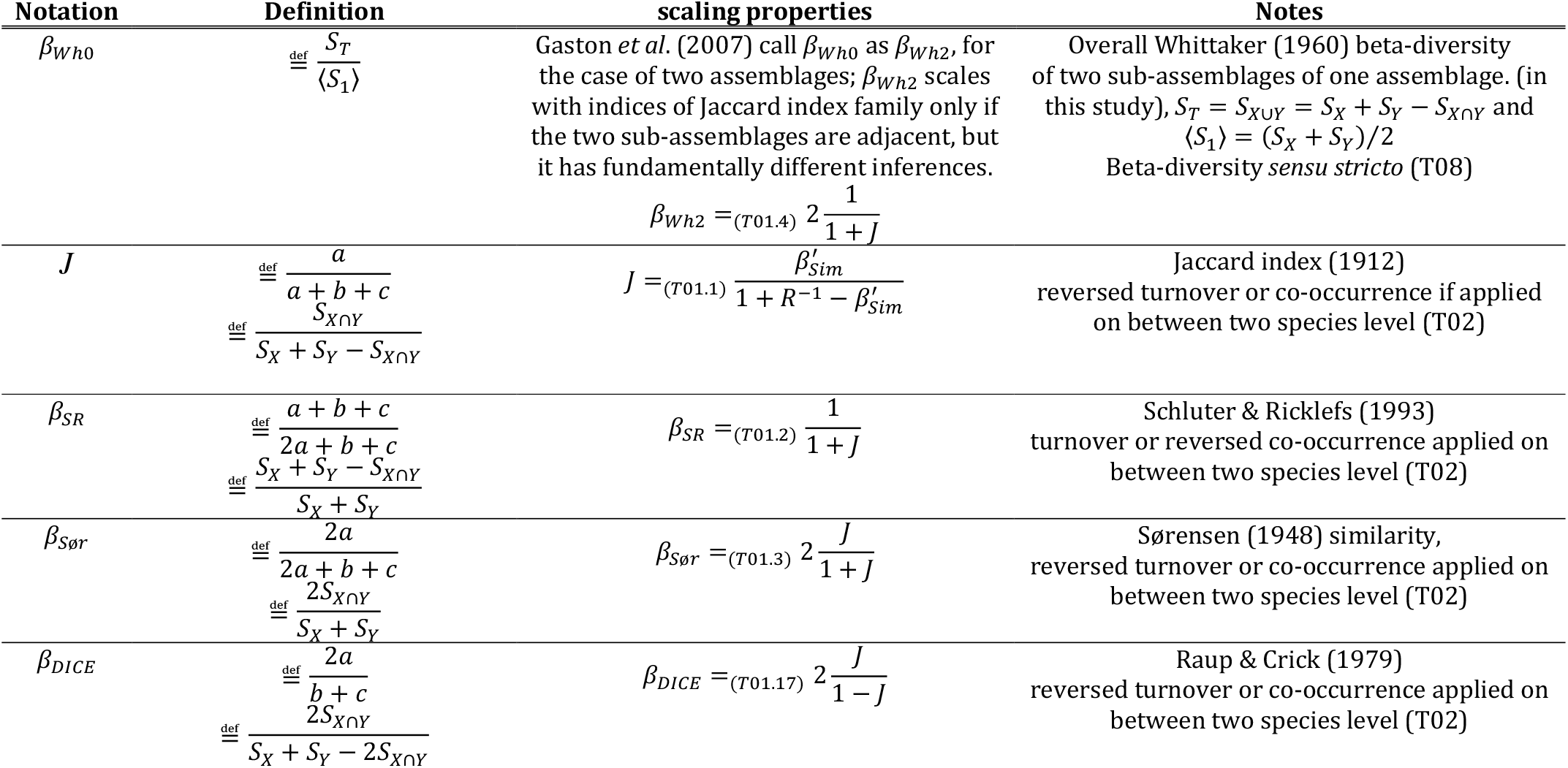
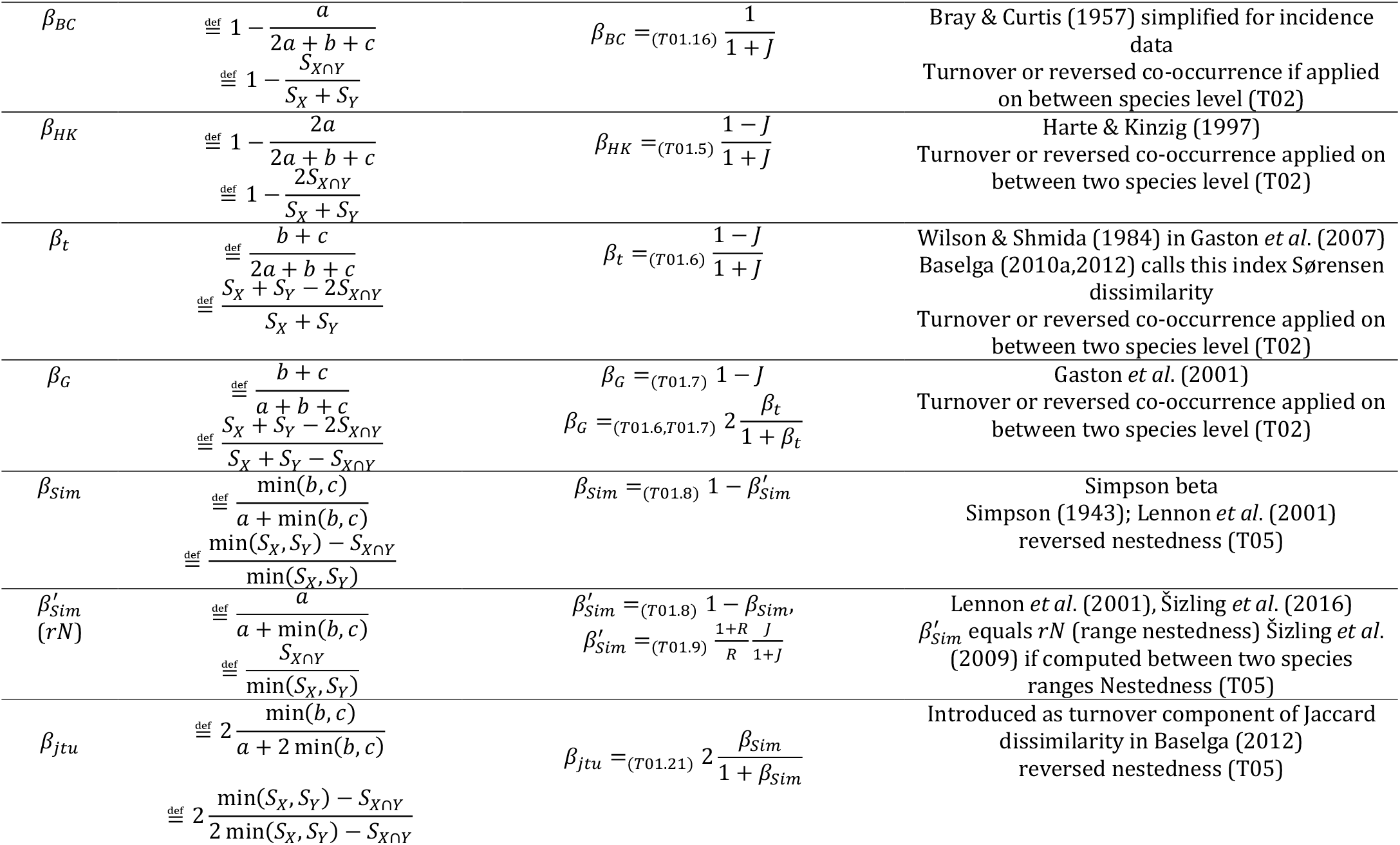
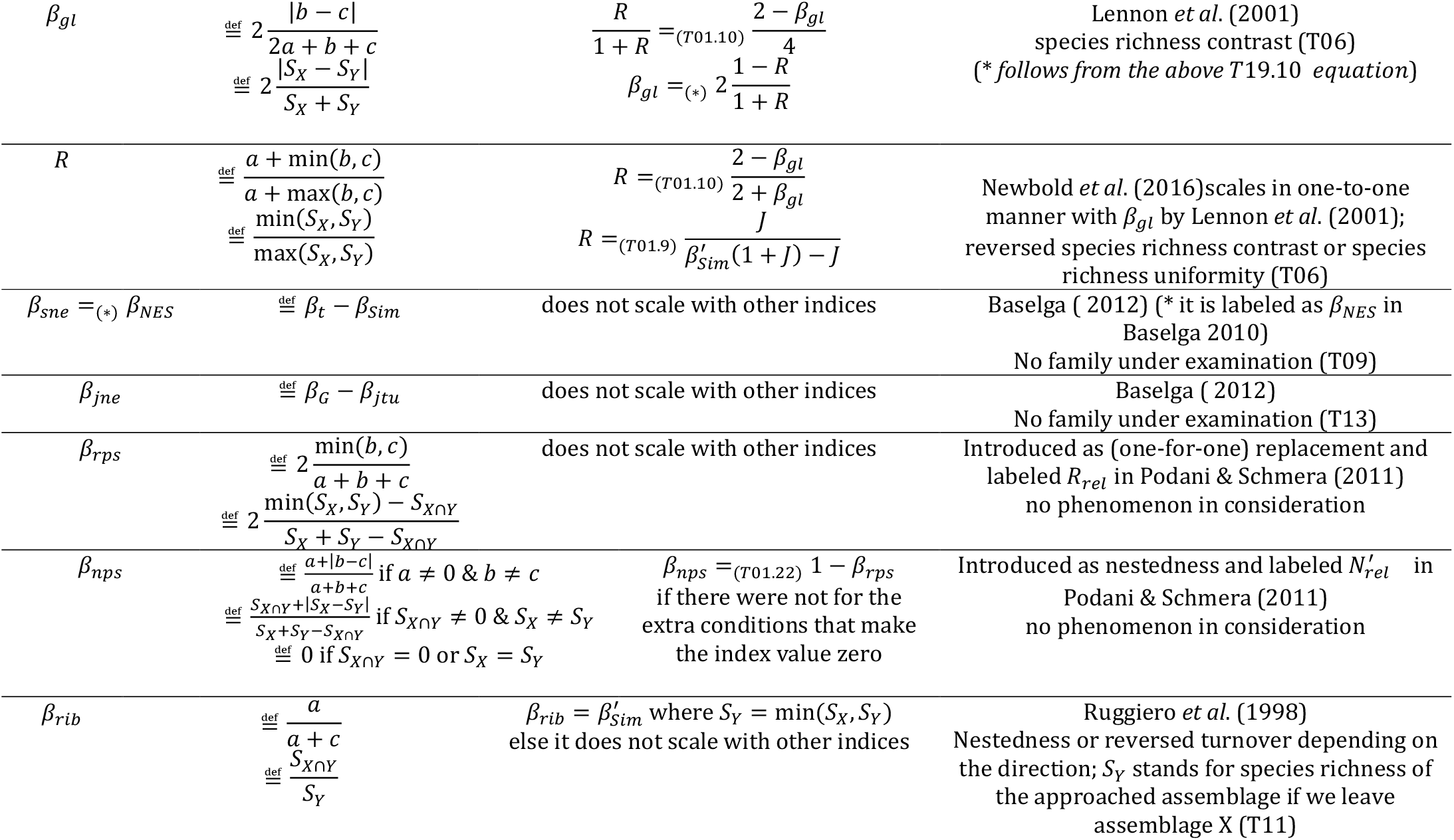
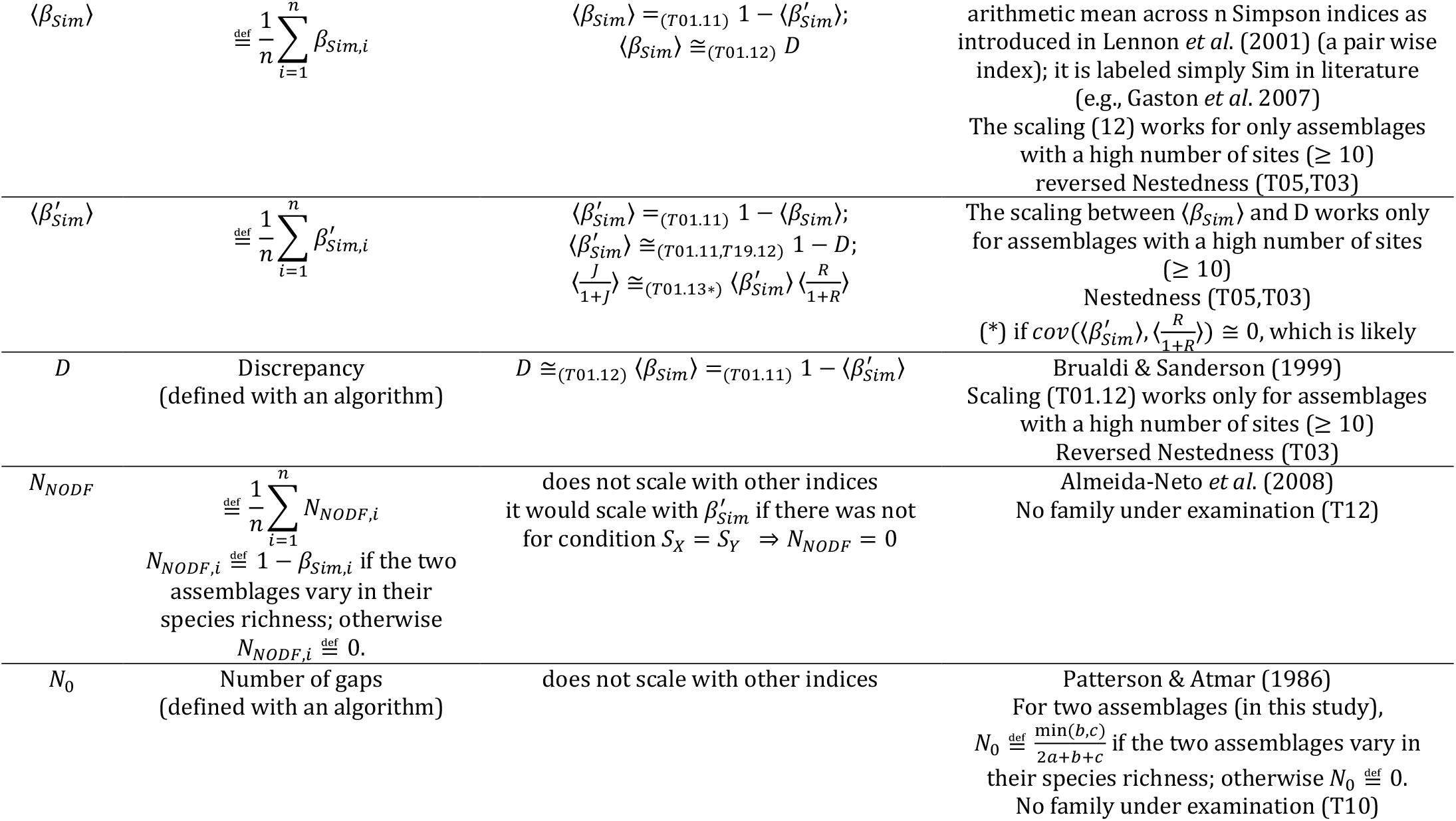
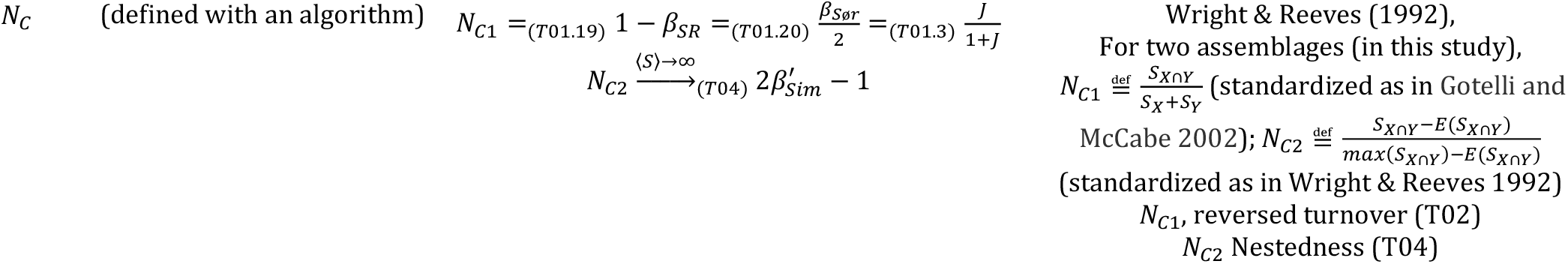
(definitions and scaling properties) The table shows the definitions of diversity indices and their mutual scaling properties. The references (e.g., T01.4) attributed to the equation marks refer to the derivations of the relationships (see Theses and Proofs T01 for derivations). The equation marks without references are derived through simple rearrangements of the equations listed in the table. All listed scaling properties are exact except those marked as ‘ ≅ ‘. Where possible, indices are defined according to Koleff *et al*. (2003) and Gaston *et al*. (2007); see the column ‘Notes’ for exceptions. Consistent with Koleff *et al*. (2003), and Gaston *et al*. (2007), *a* refers to the number of species shared by the focal assemblages 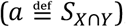, and *b* and *c* represent the numbers of species unique to the first and second assemblages (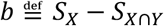 and 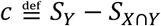). Further references can be found in the ‘Notes’ column.

### Theses (theses and proofs)

Theses T1-21 state whether or not each index from Table R1 satisfies the constraints of the spatial phenomena (Figure 3 in Šizling et al. 202X), the conditions for i-independnce of the indices, and the scaling between indices within a family and between families. The evidences for the theses employ three parameters: *a*, (the number of shared species, *S*_*X*∩*Y*_,), *b* (the number of species exclusive to the first assemblage, *S*_*X*_ − *S*_*X*∩*Y*_), and *c* (the number of species exclusive to the second assemblage, *S*_*Y*_ − *S*_*X*∩*Y*_). The individual arrangements in Figure 3 in Šizling et al. (202X), are characterized as follows: *a* = 0 for the arrangements r3 and r4; *b* = 0 in arrangements r1,r2,r5 and r6; *b* = 0 and *c* = 0 in arrangement r5; and by *c*_*r*6_ < *c*_*r*2_ < *c*_*r*1_ in arrangements r1,r2,r6.

Understanding the evidence for the theses requires a basic knowledge of linear algebra.

Specifically, one should know: what a system of linear equations is and how it can be converted to a matrix (e.g., system: *a*_11_*x*_1_ + *a*_12_*x*_2_ = *b*_1_ ; *a*_21_*x*_1_ + *a*_22_*x*_2_ = *b*_2_; the matrix: 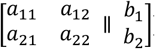. the Gauss elimination method; Cramer’s rule and the theorem stating that a nonzero determinant indicates a set of mutually independent (in our work, i-independent) equations while a zero determinant indicates a set of mutually (i-)dependent equations. Formore details on the link between linear algebra and indices of diversity see Box 2 and Box 3 in Šizling et al. (202X).

**T01** (scaling properties): *Relationships between the focal indices obey the equations listed in Table R1*. The evidence for the relationships is as follows (the numbering ‘1-22’ refers to the labels associated with equation marks in Table *R1*; brackets ⟨. ⟩ denote a mean value):

1. 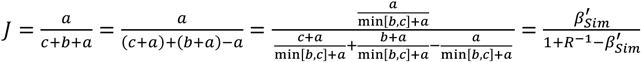.
2. 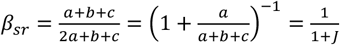.
3. 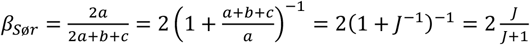.
4. 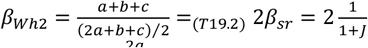.
5. 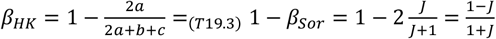.
6. 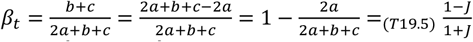.
7. 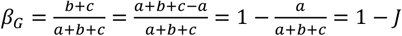.
8. 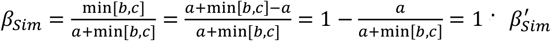.
9. 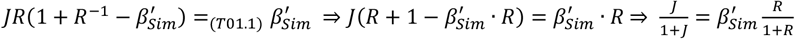.
10. Let *b* ≤ *c* then 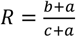 and thus 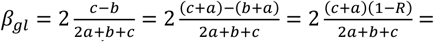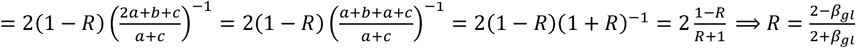.
11. 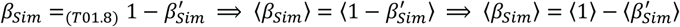
12. Simulation based evidence (see Figure R2 below).
13. 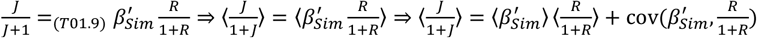. Where the i-th values 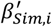 and 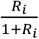 are s-independent across all i,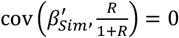. Nestedness, 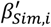 and species richness uniformity 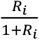 does not constraint each other (share only one sr component of the *S*_*X*_, *S*_*Y*_, *S*_*X*∩*Y*_) and are likely s-independent (implying zero covariance). However, an ecological driver makes covariance nonzero and then the relationship in Table R1 does not work.
14. 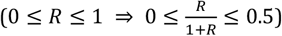, it follows 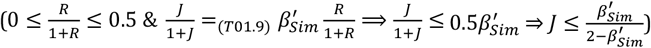.
15. 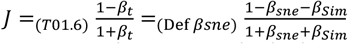.
16. 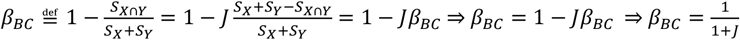.
17. 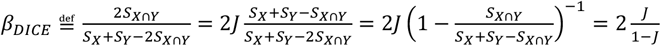.
18. 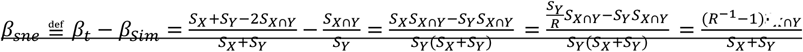. *S*_*Y*_ ≤ *S*_*X*_ is an arbitrary choice without losing generality.
19. 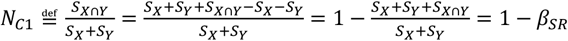.
20. 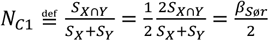.
21. 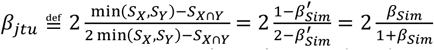.
22. 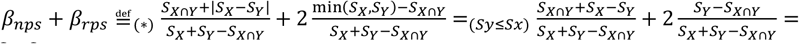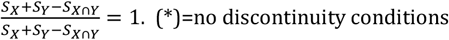.

**T02**: *Jaccard index* (*J*) *is consistent with the constraints on reversed turnover*.

**Note**: All the indices that follow strictly increasing function of *J* are consistent with the constraints on reversed turnover, and all indices that follow a strictly decreasing function of *J* are consistent with the constraints on turnover. For the scaling, see Fig. 5 in Šizling et al. (202X), Thesis T01, and Table R1. For a new index, consult Repository RI4: Calculator.

**Evidence:** 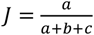. Here *a* = 0 in re-arrangements r3 and r4 in Figure 3 in Šizling et al. (202X). Hence *J*[*r*3] = *J*[*r*4] = 0, which excludes the species richness contrast but supports turnover as defined in our framework. Additionally, *J*[*r*1] < *J*[*r*2] < *J*[*r*6] < *J*[*r*5] in Figure 3 in Šizling et al. (202X). Hence, 0 < *J*[*r*1] < *J*[*r*5] = 1, which supports only the reversed constraints of turnover.

**T03:** *Discrepancy* (*D*) *for a high number of sites approaches a function that is consistent with the constraints on nestedness*.

**Note:** Figure R2 shows that more than 10 sites are sufficient to achieve consistency with the constraints.

**Evidence:** For a small number of sites to compare, index *D* does not follow any of the focal phenomena. This evidence is based on *D* computed for various pairs of sets (M1 = M2 = 15, M3=10, M4 = M5 = M6 = M7 = M8 = M9 = M10 =M11=5, and M12=4.). The rough (non-standardized) Discrepancy is defined as the minimum number of incidences that must be shifted along rows of an incidence matrix (rows represent sites; columns represent species) to achieve absolutely nested assemblages (Brualdi and Sanderson 1999). This definition simplifies to ‘min(*S*_*X*_, *S*_*Y*_) − *S*_*X*∩*Y*_’ for two assemblages. Index D is standardized by the number of incidences within the focal matrix, thus *D* = min(*S*_*X*_, *S*_*Y*_) − *S*_*X*∩*Y*_⁄(*S*_*X*_ + *S*_*Y*_) in the case. The computed order is therefore 0 = *D*[*r*1] = *D*[*r*2] = *D*[*r*5] = *D*[*r*6] < *D*[*r*3] < *D*[*r*4] = 0.5. The equality *D*[*r*1] = *D*[*r*5] excludes species richness contrast, nestedness, and turnover, and the inequality *D*[*r*3] < *D*[*r*4] excludes nestedness, but not contrast. However, if the number of sets is large enough (simulations suggest more than 30 simulations), the standardization by the number of incidences begins to work properly and the index *D* will scale with the indices of nestedness (Figure R2 below). We demonstrate the reason using two extreme cases: with maximum and minimum possible D. The Discrepancy of an absolutely nested matrix is by definition zero (no shift of incidences is needed). Maximum Discrepancy occurs in a matrix where almost all incidences (except the incidences of one site) must be shifted to gain an absolutely nested matrix. The Discrepancy of an absolutely non-nested matrix (where each site has its unique set of species) is then computed as the total species richness (sum across all sites) of species that has to be shifted (i.e., excluding one of the most species rich sites). This equals 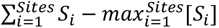, where *S*_*i*_ is the species richness of the i-th site. It is standardized by the number of incidencies, i.e.,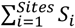. Hence, 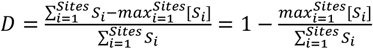, which approaches one if 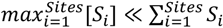. This is the case of practically all datasets with large numbers of sites, and therefore *D* ≅ 1 can be attributed to absolutely non-nested multisite assemblages. It is apparent that all matrices between these two extremes have D values between 0 and 1. The condition 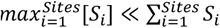 might only be violated if the maximum species richness was high and species richness of the other sites was extremely small, which is unlikely. Simulations show a one-to-one scaling of *D* with Simpson beta (non nestedness) for matrices of 30 and 100 sites (Figure R2).

**T04:** *N*_*C*_, s*tandardized as in* Wright & Reeves (1992) *(here labeled N*_*C*2_; *labeled as C in Wright & Reeves 1992) approaches i-dependence on Simpson beta as Species Richness approaches infinity. N*_*C*2_ *is i-dependent on species richness*.

**Note:** N_C2_ is thus consistent with the constraints on Nestedness.

**Evidence:**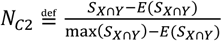, where max(*S*_*X*⋂*Y*_) = *S*_*Y*_ if we put *S*_*X*_ ≥ *S*_*Y*_, and E() stands for expectance. So, 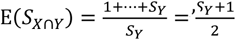. Hence 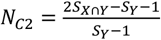. In general, 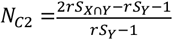, where *r* > 0 emulates variation in species richness. Apparently, *N*_*C*2_ is idependent on r, and 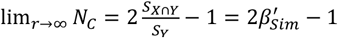.

**T05:** *Simpson beta β*_*Sim*_ and *β*_*jtu*_ *are consistent with the constraints on reversed nestedness and* 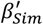 *is consistent with the constraints on nestedness*.

**Note:** The evidence is done for *β*_*Sim*_. For the strictly increasing function of *β*_*jtu*_ with *β*_*Sim*_, and the strictly decreasing function of 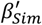 with *β*_*Sim*_ see Fig. 5 in Šizling et al. (202X), Thesis T01, and Table R1.

**Evidence:** 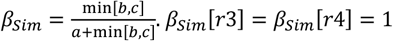 because *a* = 0 in these cases; *β*_*Sim*_[*r*1] = *β*_*Sim*_[*r*2] = *β*_*Sim*_[*r*5] = *β*_*Sim*_[*r*6] = 0 because one of the variables *b, c* equals zero and *a* ≠ 0 in these cases. All the indices are continuous and none of them depends on max(*b, c*). For scaling between 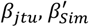 and *β*_*Sim*_ see (T01.8, T01.21 and Figure 5 in Šizling et al. 202X).

**T06:** *β*_*gl*_ *and R are consistent with the constraints on species richness contrast and species richness uniformity (i*.*e*., *reversed species richness contrast), respectively*.

**Evidence:** R is the ratio of minimum to maximum species richness. Hence *R*[*r*1] = *R*[*r*3] < *R*[*r*2] < *R*[*r*6] < *R*[*r*4] = *R*[*r*5] = 1. *R* is therefore inversely related to species richness contrast, representing species richness uniformity. *β*_*gl*_ scales negatively with *R*, (see T01.10).

**T07:** *β*_*nps*_ *is inconsistent with the constraints on nestedness, turnover or species richness contrast*.

**Evidence:** If we ignore the extra conditions that make *β*_*nps*_ discontinuous (i.e., *β*_*nps*_ if *S*_*X*∩*Y*_ = 0 or *S*_*X*_ = *S*_*Y*_) then 0 = *β*_*nps*_[*r*4] < *β*_*nps*_[*r*3] < *β*_*nps*_[*r*5] = *β*_*nps*_[*r*6] = *β*_*nps*_[*r*2] = *β*_*nps*_[*r*1] = 1, where r1,…,r6 are re-arrangements of two assemblages defined in Figure 3 in Šizling et al. (202X). The inequality *β*_*nps*_[*r*4] < *β*_*nps*_[*r*3] excludes nestedness, the equality *β*_*nps*_[*r*2] = *β*_*nps*_[*r*1] excludes the species richness contrast, and the equality *β*_*nps*_[*r*2] = *β*_*nps*_[*r*1] excludes turnover.

If we add the condition: “*β*_*nps*_ = 0 if *S*_*X*∩*Y*_ = 0” (the variant of the index under discussion in Šizling et al. 202X, which is labeled *N*_*rel*_ in Podani and Schmera 2011), then the relations become 0 = *β*_*nps*_[*r*4] = *β*_*nps*_[*r*3] < *β*_*nps*_[*r*5] = *β*_*nps*_[*r*6] = *β*_*nps*_[*r*2] = *β*_*nps*_[*r*1] = 1. This again excludes species richness contrast and turnover, but it agrees with the constraints on nestedness (Figure 3 in Šizling et al. 202X). This is only due to the extra condition, so if we consider that indices must follow the requirement of continuity, the *β*_*nps*_ does not capture the nestedness. However, there is another reason why *β*_*nps*_does not capture the experience of nestedness. The constraints *N*[*r*1] = *N*[*r*2] implies independence of nestedness from the species richness of the richer assemblage. If we violate this condition where there is an overlap between the assemblages (i.e., *S*_*X*∩*Y*_ > 0), then we are inconsistent with the constraint *N*[*r*1] = *N*[*r*2] (Figure 3 in Šizling et al. 202X). The intuition is that if we enter the sea, then the fraction of how much we are immersed does not depend on the size of the sea. It is easy to see that 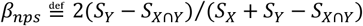 changes with changing *S*_*X*_ (= max(*S*_*X*_, *S*_*Y*_)), which excludes nestedness (arbitrarily without loss of universality *S*_*Y*_ ≤ *S*_*X*_).

If we add the condition: “*β*_*nps*_ = 0 if *S*_*X*∩*Y*_ = 0 or *S*_*X*_ = *S*_*Y*_” (this variant of the index that is labeled 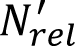 in Podani and Schmera 2011) then the relations turn into 0 = *β*_*nps*_[*r*4] = *β*_*nps*_[*r*3] = *β*_*nps*_[*r*5] = *β*_*nps*_[*r*6] = *β*_*nps*_[*r*2] = *β*_*nps*_[*r*1] = 1. Again, the equality *β*_*nps*_[*r*3] = *β*_*nps*_[*r*5] excludes nestedness, the equality *β*_*nps*_[*r*2] = *β*_*nps*_[*r*1] excludes the species richness contrast, and the equality *β*_*nps*_[*r*2] = *β*_*nps*_[*r*1] excludes turnover.

**T08:** Index *β*_*rps*_ *is inconsistent with the constraints on nestedness, turnover and species richness contrast. The index β*_*rps*_ *is also inconsistent with the concept of One-for-One Replacement*.

**Evidence:** The relations for *β*_*rps*_ are reversed to the relations for *β*_*nps*_ where no extra conditions that make the index discontinuous are considered (i.e., 0 = *β*_*rps*_[*r*4] < *β*_*rps*_[*r*3] < *β*_*rps*_[*r*5] = *β*_*rps*_[*r*6] = *β*_*rps*_[*r*2] = *β*_*rps*_[*r*1] = 1), for *β*_*rps*_ = 1 − *β*_*nps*_ in this case (T01.22). Therefore, the reasoning for nestedness, species-richness contrast, and turnover follows the evidence T07 above. In the case of one-for-one replacement, there is a constraint *R*[*r*4] = *R*[*r*3] on the one-for-one replacement that follows the point of Podani & Schmera (2011). The inequality *β*_*rps*_[*r*4] < *β*_*rps*_[*r*3] therefore excludes one-for one replacement. Because the numerator of *β*_*rps*_ for r3 equals the numerator of *β*_*rps*_ for r4, the problem is introduced by denominator (i.e., standardization of the index with *S*_*X*_ + *S*_*Y*_ − *S*_*X*∩*Y*_). One may argue that the one-for-one replacement of r3 differs from one-for-one replacement of r4, and thus the inequality *β*_*rps*_[*r*4] < *β*_*rps*_[*r*3] does not disqualify *β*_*rps*_ as a measure of the one-for-one replacement. In this case, however, there should be a difference between *β*_*nps*_(*S*_*X*_ = 8, *S*_*Y*_ = 20, *S*_*X*∩*Y*_ = 1) and *β*_*nps*_(17,20,10), but *β*_*nps*_(8,20,1) = *β*_*nps*_(17,20,10) (Figure 1 in Šizling et al. 202X).

**T09:** *Nestedness by Baselga* (*β*_*sne*_) *is inconsistent with the constraints on nestedness, turnover and species richness contrast*.

**Evidence:** 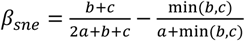. For re-arrangements r1 and r2 in Figure 3 in Šizling et al. (202X), *β*_*sne*_[*r*1] < *β*_*sne*_[*r*2], because *β*_*Sim*_[*r*1] = *β*_*Sim*_[*r*2] = 0, *b* = 0, *c*_*r*2_ < *c*_*r*1_, and *a* does not vary between these two cases. This excludes nestedness. *β*_*sne*_[*r*5] = *β*_*sne*_[*r*3] = 0, because *b* = *c*, which excludes turnover. *β*_*sne*_[*r*1] > *β*_*sne*_[*r*3] = 0, which excludes species richness contrast (*β*_*sne*_[*r*3] = 0, because *a* = 0; and (*β*_*sne*_[*r*1] > 0, because *min*(*b, c*) = 0 and *b* ≠ *c*).

**T10:** *Number of gaps* (*N*_0_) *is inconsistent with the constraints on nestedness, turnover and species richness contrast*.

**Evidence:** This evidence is based on *N*_0_ indices computed for various pairs of sets (M1 = M2 = 15, M3=10, M4 = M5 = M6 = M7 = M8 = M9 = M10 =M11=5, and M12=4.). The computed order is 0 = *N*_0_[*r*1] = *N*_0_[*r*2] = *N*_0_[*r*4] = *N*_0_[*r*5] = *N*_0_[*r*6] < *N*_0_[*r*3] = 0.3 for the index that was standardized by number of incidences within the focal matrix. This matches no experience of the five spatial phenomena.

**T11:** *The Ruggiero index of beta-diversity* (*β*_*Rib*_) *is consistent with the constraints on reversed turnover or nestedness depending on the direction*.

**Evidence:** The index *rib* depends on the order of the focal assemblages. It is defined as 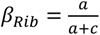, where *c* captures either the first or second assemblage. In our case, 0 = *c*_*r*5_ < *c*_*r*6_ < *c*_*r*2_ < *c*_*r*1_ and *a*_*r*5_ = *a*_*r*6_ = *a*_*r*2_ = *a*_*r*1_. Hence, 0 < *β*_*Rib*_[*r*1] < *β*_*Rib*_[*r*5]. At the same time 0 = *β*_*Rib*_[*r*3] < *β*_*Rib*_[*r*4], because *a* = 0 in these cases. The index *β*_*Rib*_ thus captures reversed turnover. If we replace *b* with *c*, then *β*_*Rib*_[*r*1] = *β*_*Rib*_[*r*2] = *β*_*Rib*_[*r*6] = *β*_*Rib*_[*r*5] = 1, because *c*=0 in these cases. The index *rib* therefore captures nestedness in the case.

**T12:** *Nestedness by Almeida-Neto et al. (2008)* (*N*_*NODF*_) *would be consistent with the constraints on nestedness if we ignored the condition that S*_*X*_ = *S*_*Y*_ ⟹ *N*_*NODF*_= 0.

**Evidence:** Where assemblages differ in their species richness, *N*_*NODF*_ equals 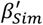 from the Simpson beta family, which is an index of nestedness. Where assemblages have equal species richness *N*_*NODF*_ = 0. If we accepted that two equal sized assemblages are not mutually nested (*NODF* (*r*5) = 0, but *NODF* (*r*6) = 1), then *N*_*NODF*_ would be an index of nestedness. However, our framework excludes this possibility.

**T13:** *The Nestedness resultant component of Jaccard dissimilarity (β*_*jne*_*) is inconsistent with the constraints on nestedness, turnover or species richness contrast*.

**Evidence:** This evidence is based on *N*_0_ indices computed for various pairs of sets (M1 = M2 = 15, M3=10, M4 = M5 = M6 = M7 = M8 = M9 = M10 =M11=5, and M12=4.). The computed order is *β*_*jne*_[*r*3] = *β*_*jne*_[*r*4] = *β*_*jne*_[*r*5] = 0, *β*_*jne*_[*r*6] = 0.2, *β*_*jne*_[*r*2] = 0.5, and *β*_*jne*_[*r*1] ≅ 0.7. The equality *β*_*jne*_[*r*4] = *β*_*jne*_[*r*5] excludes nestedness and turnover, and tha equality *β*_*jne*_[*r*3] = *β*_*jne*_[*r*4] excludes species richness contrast.

**T14:** *An algorithm to compute Jaccard similarity and Simpson nestedness from two i-independent indices if one of the indices is partitioned*.

**The algorithm:** Let

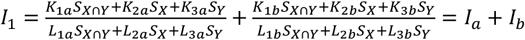

(*if index I*_*b*_ *is subtracted the k*_*ib*_ *coefficients are multiplied by ‘*−1*’*) and

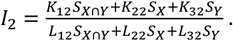

Then the triple *I*_*a*_, *I*_*b*_, and *I*_2_ are necessarily i-dependent. Thus

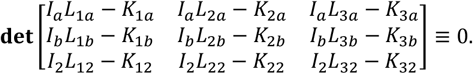

*I*_2_, and *I*_1_ are known (*I*_1_ = *I*_*a*_ + *I*_*b*_), so

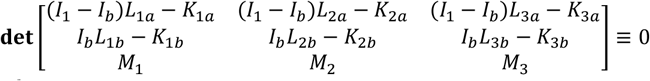

where *M*_*i*_ = *I*_2_ *L*_*i*2_ − *K*_*i*2_.

After expansion

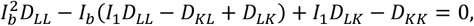

where 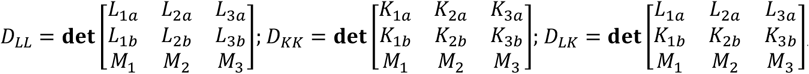 and 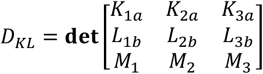.

Having *I*_*b*_, we can pick up two i-independent indices and then use the algorithm from Box 3 in Šizling et al. XXXX to compute any dimensionless index.

This solution works only if the indices *I*_*a*_, *I*_*b*_ are mutually i-independent, the indices *I*_*a*_, *I*_1_ are mutually i-independent, and the indices *I*_1_, *I*_*b*_ are mutually i-independent (together they are always i-dependent). If any pair of indices is mutually i-dependent, the solution is simpler: convert the i-dependent indices to an index from their family (Table R1) and then follow the algorithm from Box 3 in Šizling et al. XXXX.

**T15:** Symmetric and linear indices with equal denominators are i-dependent.

That is: Let us have two indices that are defined as

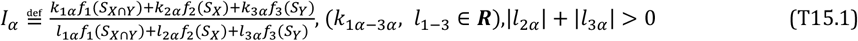

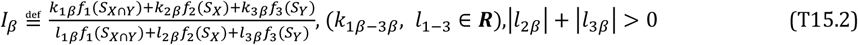

where *f*_*i*_(*X*) are strictly increasing or strictly decreasing functions from ***R***^+^ to ***R***^+^, and *k*_2*α*_= *k*_3_*α*, *k*_2*β*_= *k*_3 *β*_, and *l*_1_*α* = *l*_2*α*_, *l*_1 *β*_= *l*_2*β*_, then (*l*_1 *α*_= *l*_1 *β*_, & *l*_12*α*_= *l*_2*β*_,) ⇒ *I*_*α*_ is i-dependent on *I*_*β*_.

**Note 1:** if *f*_*i*_(*X*) = *X*, ∀*X* ∈ ***R***, then Eqs. T15.1 and T15.2 turn into Eq. 8 in Šizling et al. 202X.

**Note 2:** It follows that two asymmetric indices with equal denominators are i-independent.

**Note 3:** The symmetry required here is a stronger form of symmetry as it necessitates symmetry in both the numerator and denominator separately. For example, the index R is not a symmetric index according to this theorem.

**Evidence:**

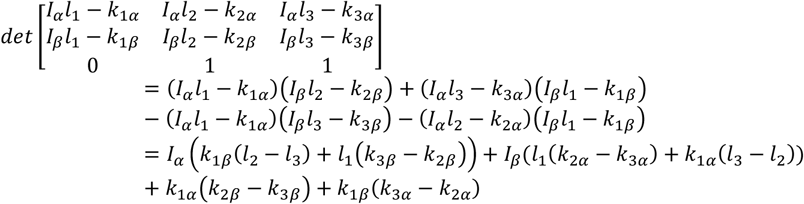

Therefore determinant is zero (i.e., indices are i-dependent) If *l*_2_ − *l*_3_ = 0 & *k*_3 *α*_ − *k*_2*α*_= 0 & *k*_3 *β*_ − *k*_2*β*_= 0.

**T16:** (*species rich and poor assemblages*): The variation of *S*_*poor*_ = 2⟨*S*⟩*R*/(1 + *R*) indicates inevitable change in richness of the species poorer assemblage, and the variation of *SR*_*rich*_ = 2⟨*S*⟩/(1 + *R*) indicates inevitable change in the richness of the species richer assemblage.

**Evidence:** The evidence is based on the solution of the system of three equations for 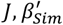 and ⟨*S*⟩. See Box 2 in Šizling et al. 202X. If *S*_*X*_, *S*_*Y*_, and *S*_*X*∩*Y*_are mutually i-independent, the system follows 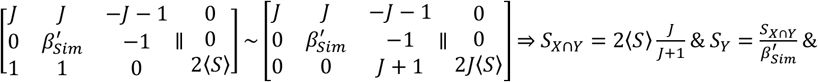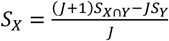. It follows that 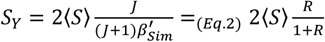. The last equality follows from Eq. 2. Finally,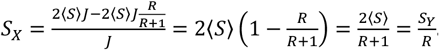. Because 0 < *R* ≤ 1, then *S*_*Y*_ ≤ *S*_*X*_, and we relabel *S*_*poor*_ ≔ *S*_*Y*_ and *SR*_*rich*_ ≔ *S*_*X*_. If *S*_*X*_, *S*_*Y*_, and *S*_*X*∩*Y*_are mutually i-dependent, then 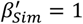 or *R* = 1 and the derived equations for *S*_*poor*_ and *SR*_*rich*_ remain valid.

**T17:** *Jaccard index (J) is consistent with conceptual predefinition of co-occurrence if applied to more sites occupied by two species instead of more species that belong to two assemblages*.

**Evidence:** If we consider the Venn diagrams in Figure 3 (Šizling et al. 202X) to show two species ranges (sets of occupied sites), then the constraints on Co-occurrence follow:

Min = Co[r4] = Co[r3] < Co[r1] < Co[r2] < Co[r6] < Co[r5] = Max.

Because 0 = *J*[r4] = *J*[r3] < *J*[r1] < *J*[r2] < *J*[r6] < *J*[r5] = 1, the evidence is completed.

**T18:** The system of Eq. 10 and Eq. 11 in Šizling et al. (202X) is mutually i-independent if *S*_*X*∩*Y*_ ≠ 0 and *S*_*X*_ ≢ *S*_*Y*_ ≢; *S*_*X*∩*Y*_ ≢ *S*_*X*_.

**Evidence:** If *S*_*X*_, *S*_*Y*_, and *S*_*X*∩*Y*_are mutually i-independent, then the system follows 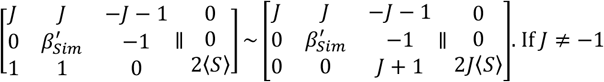 and *J* ≠ *0* and ⟨ *S* ⟩ ≠ 0 (mean S), this matrix provides a unique and non-trivial (i.e., non-zero) solution of the system. Alternatively 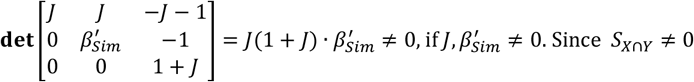 implies 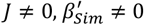, and ⟨*S*⟩ ≠ 0, it follows that 2*J*⟨*S*⟩ ≠ 0 and the solution is not trivial (it is non zero). As an alternative, the last scaling equation can be replaced with *S*_*X*_ + *S*_*Y*_ − *S*_*X*∩*Y*_ = *S*_*TOT*_ where *S*_*TOT*_ is species richness of both sites. If the condition ‘*S*_*X*∩*Y*_ ≠ 0 and *S*_*X*_ ≢ *S*_*Y*_ ≢ *S*_*X*∩*Y*_ ≢ *S*_*X*_’ is violated, then each index definition is a linear function of two variables and thus two equations are enough to get a unique solution, making one of the equations i-dependent on the others.

**T19:** *(i-dependence of three indices): If S*_*X*∩*Y*_ ≠ 0 *(i*.*e*., *J* ≠ 0*), then the value of any index defined by Eq. 8 in Šizling et al. (202X) can be computed from values of Jaccard index (J) and Simpson nestedness* 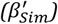 *using*

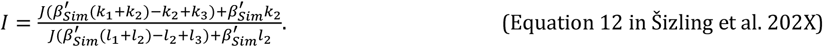

**Evidence:** Consider arbitrary *S*_*Y*_ ≤ *S*_*X*_. Then *S*_*X*∩*Y*_ ≠ 0 ⇒ *S*_*Y*_ ≠ 0 ⇒ *S*_*X*_ ≠ 0. The new index is defined as 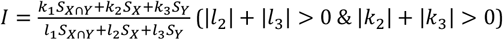 which is the definition Eq. 8 in Šizling et al. (202X). The Eq. 12 results as a solution of the determinant equation 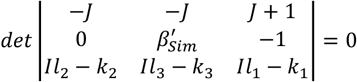. The proof of the i-independence follows the reverse logic. (i) substitute definitions of *J* and 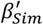 into Eq. 12, (ii) convert Eq. 12 into the third equation of the system of linear equations and (iii) show that the determinant equals zero. In the same way is proved the equation for triplets *I, J, R* and 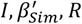. In these cases the restriction *S*_*X*∩*Y*_ ≠ 0 is replaced with *S*_*Y*_ ≠ 0.

**T20:** *If Šizling et al. (2016) published only two partitioned indices (β*_*sne*_, *β*_*nps*_*) then the referred information would be unequivocal*.

**Evidence:** When we convert the indices reported by Šizling et al. (2016) into two partitioned indices, we got *β*_*sne*_ = 0.1, *β*_*nps*_ = 0.61 for the pre-agricultural landscape, and *β*_*sne*_ = 0.00, *β*_*nps*_ = 0.51 for early agricultural landscape. These indices suggest: *S*_*X*_ ≅ 31.01, *S*_*Y*_ ≅ 22.99, *S*_*X*∩*Y*_ ≅ 15.48 or *S*_*X*_ ≅ 41.63, *S*_*Y*_ ≅ 12.37, *S*_*X*∩*Y*_ ≅ 2.28 for pre-agricultural landscape; and *S*_*X*_ ≅ 32.00, *S*_*Y*_ ≅ 32.00, *S*_*X*∩*Y*_ ≅ 21.60 or *S*_*X*_ ≅ 48.32, *S*_*Y*_ ≅ 15.68, *S*_*X*∩*Y*_ ≅ 0.00 for early agricultural landscape (thesis T14 or Repository R4: calculator). This can be interpreted as either increasing species richness uniformity (*R* increases from 0.74 to 1), or decreasing species richness uniformity (*R* decreases from 0.74 to 0.32) species richness uniformity. This dual interpretation precludes any correct conclusion.

**T21:** *Any dimensionless index of turnover (defined by the constraints in Fig. 3) must have both S*_*X*_ *and S*_*Y*_ *in its denominator*.

**Evidence**: Dimensionless indices are ratios with numerators and denominators. Accurate measurement of turnover requires that information on both assemblages *S*_*X*_, and *S*_*Y*_ be represented to capture their interaction fully. Including only one of *S*_*X*_, or *S*_*Y*_ in the denominator while having both in the numerator creates an index without an upper limit. This contradicts the required bounded nature of the index, as specified by the constraints *T*[*r*4] = *T*[*r*3] = Max in Fig. 3.

**FIGURE R2.**
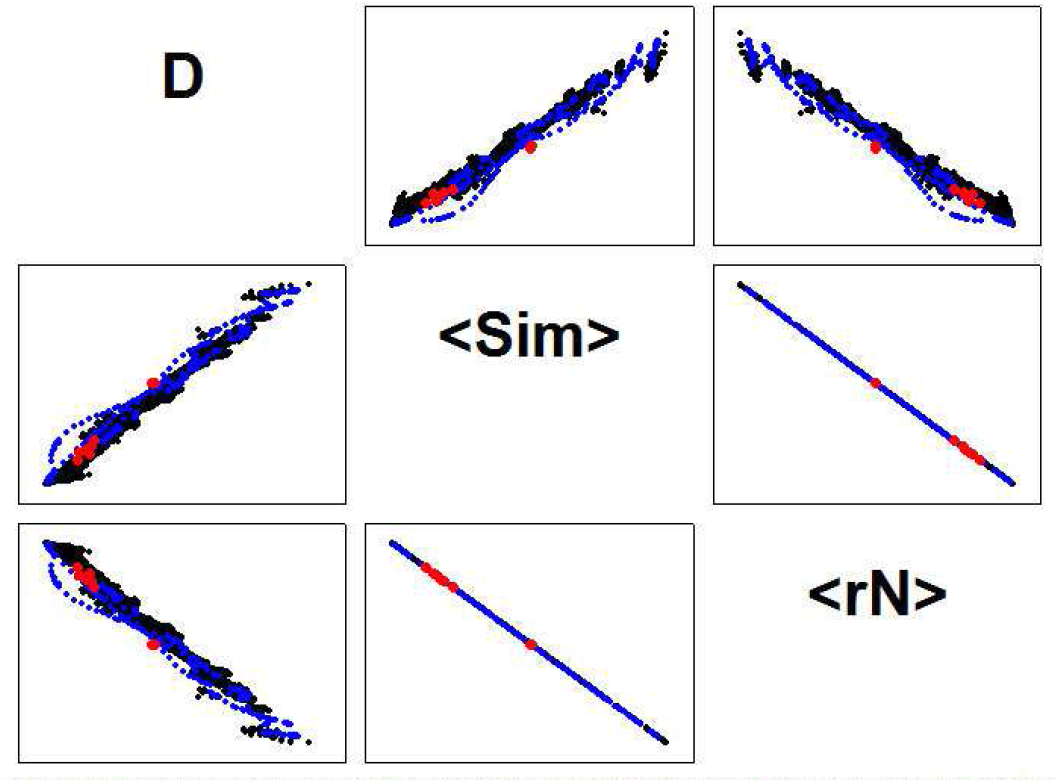
(average values) Relationships between indices of nestedness computed for more assemblages as simulated (30 sites x 30 species – black dots; 100 sites x 100 species – blue dots) and observed (red dots, R6 - data description). Matrices were generated to cover a wide spectrum of assemblages (see simulation details below). Indices include *D* (discrepancy by Brualdi & Sanderson, 1999); ⟨*Sim*⟩ (simple mean of the Simpson index across all pairs of adjacent plots; Lennon et al., 2001); and 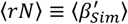 (simple mean of nestedness as defined in Šizling et al., 2009); *D*, was standardized by dividing with the total number of incidences within the focal matrix (Greve et al., 2005). The indices scale with each other in a one-to-one manner, belonging to the same family and measuring the extent of nestedness. For exact evidence, see theses T03, and T05.

**FIGURE R3.**
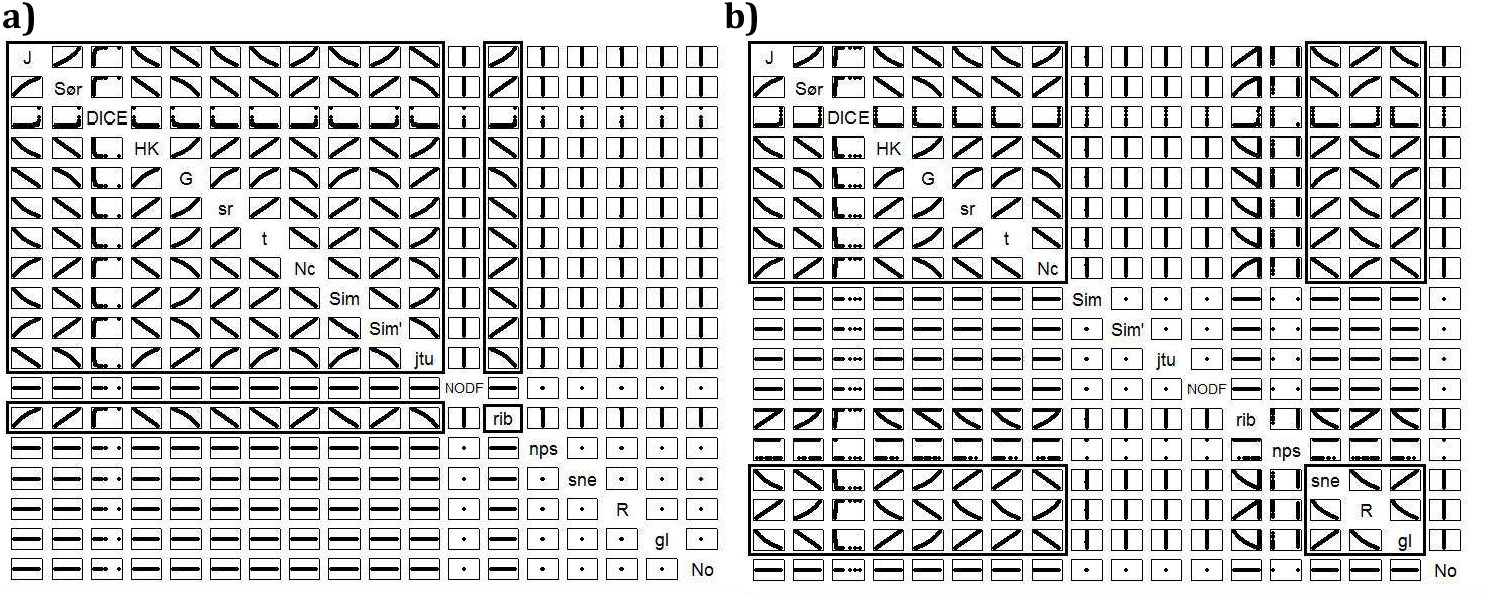
(interrelated assemblages) Relationship between pairwise indices where there is no contrast in species richness (a) and where assemblages are perfectly nested (b). As predicted, no contrast in species richness merges families of *J*, and *β*_*Sim*_ together, and perfect nestedness merges families of *J, β*_*sne*_, and *β*_*gl*_ together. The evidence that *β*_*sne*_ shows variability where nestedness is perfect disqualifies this measure from being a proxy for nestedness component. Black rectangles delimit the merged families. For a detailed legend, see capture to Figure 5 in Šizling et al. 202X.

The random points for Fig. R3 and Figs 5,6 in Šizling et al. (202X) follow *S*_*X*∩*Y*_ = *Rnd*(1) * (1 − *T*), *S*_*X*_ = *S*_*X*∩*Y*_ + *trunc*(100 * (1 − *N*) * *Rnd*(1))/100, *S*_*Y*_ = *Trunc*(100 * (*Rnd*(1) * (*min*(1, *S*_*X*_ + *C*) − *max*(0, *S*_*X*_ − *C*)) + *max*(0, *S*_*X*_ − *C*))/100. T, *N, C* ∈ ⟨0,1⟩ where T=1 indicates no shared species richness; T=0 indicates all possible overlaps in species richness; N=1 indicates absolute nestedness; N=0 indicates that nestednes is not constrained; C=1 indicates unlimited species richness contrast; C=0 indicates no species richness contrast; G=0.2 indicates a maximum 20% variation in the difference between species richness of the sites. In particular, T=0, N=0, C=1 (Figure 5 in Šizling *et al*. 202X); T=0, N=0, C=0.2 (Figure 6); T=0, N=0, C=0 (Figure R3A); T=0, N=1, C=1 (Figure R3B in Šizling *et al*. 202X).

**FIGURE R4.**
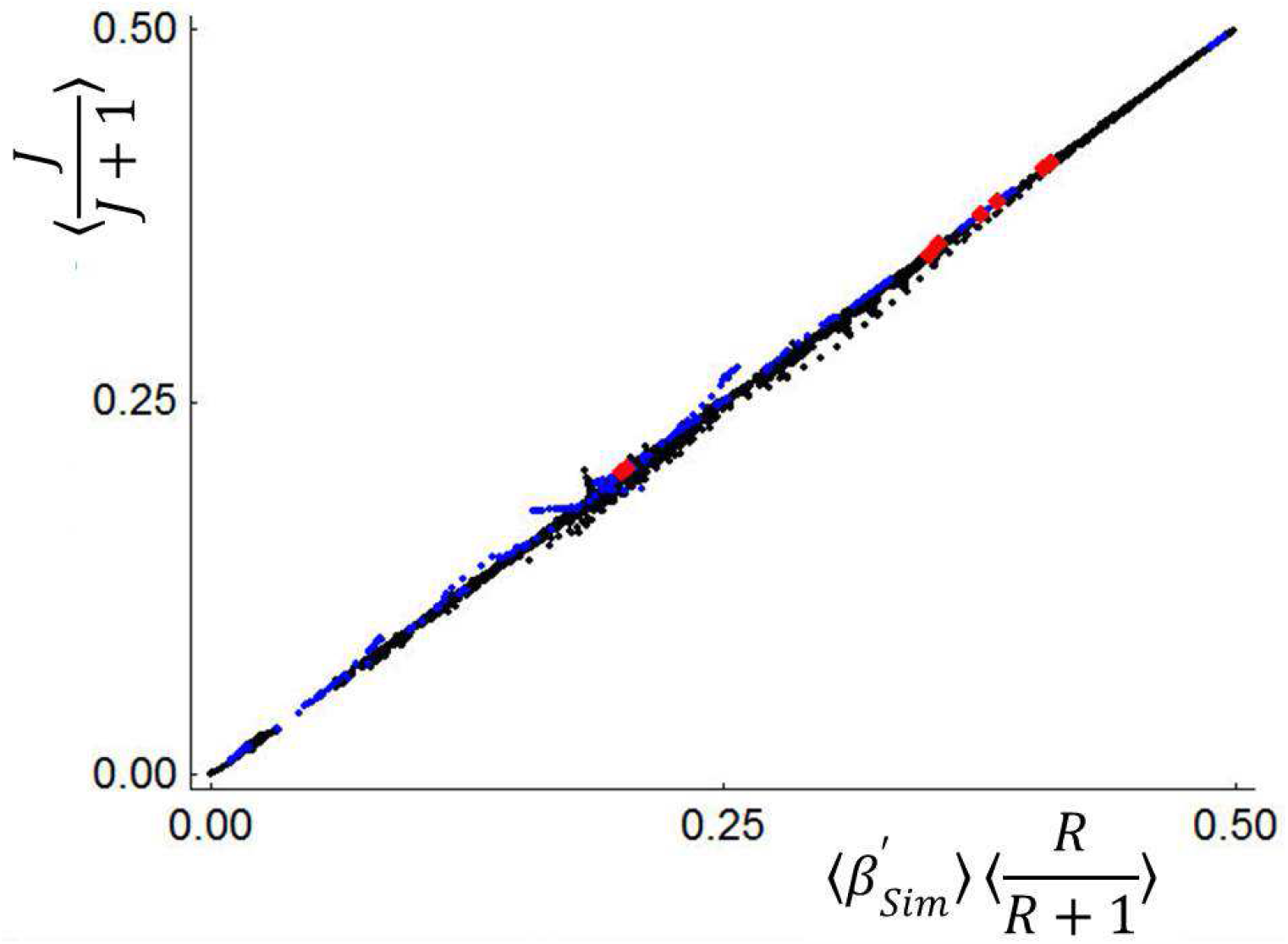
(Multisite Version of Eq. 2 in Šizling et al., 202X) The accuracy of the approximate Eq. 2 applied to mean values across multiple pairs of sites follows thesis T01.13. The exact equality between the X-values and Y values in the plot is affected by the covariance between 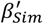 and *R*/(1 + *R*), which is expected to be nearly zero. The figure shows the extent to which this expectation is met for simulated (30 sites x 30 species – black dots; 100 sites x 100 species – blue dots) and observed (red dots) multiple-site assemblages (R6 data description and RI2: Data). The y-axis shows the left side of the equation (i.e., the mean reversed turnover; ⟨*J*/(*J* + 1)⟩), and the x-axis shows the right side of the equation (i.e., the product of the mean values of nestedness, 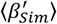 and species richness uniformity ⟨*R*/*R* + 1⟩). The points approach an identity line, showing that the covariance approaches zero as expected. The 30x30 and 100x100 assemblages were produced by the algorithm adopted from Šizling et al. (2009).

## R5 (i-independence of and s-dependence on species richness)

The Jaccard index (*J*), and thus all the indices that scale one-to-one with *J* (Figure R5), has been referred to as ‘dependent’ on species richness (Simpson, 1943; Lennon *et al*., 2001; Koleff & Gaston, 2002; Baselga, 2010a) and on the contrast between species richness of two assemblages (Simpson, 1943). This has led to a search for an index that is species richness ‘independent’, and to attempts to modify *J*, so that ecologists could compare assemblages that varied in species richness. This s-dependence between *J* and species richness was based on empirical experience (Koleff & Gaston, 2002), and on arguments that there are bounds on the *J* imposed by contrast in species richness (Simpson, 1943).

We found that all dimensionless (unitless) indices, including *J*, are i-independent of species richness. For all indices that can be expressed by the universal definition (Eq. 8 in Šizling et al. 202X) it holds that

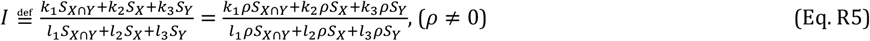

and thus the index does not change when *S*_*X*_, *S*_*Y*_, and *S*_*X*∩*Y*_ scale proportionally to each other. In this case, the index has the same value regardless of species richness, and thus it is not uniquely determined by species richness. Any observed s-dependence between the dimensionless index and species richness is therefore caused by disproportional scaling between *S*_*X*_, *S*_*Y*_ or *S*_*X*∩*Y*_.

Disproportional scaling can, however, appear at sites with small species richness because species richness is an integer. In this case, the frequency distribution of possible *J* values is affected by total species richness. This in turn affects the most likely value of *J*, imposing its s-dependence on species richness. The reason is that the Jaccard index can only have a finite number of values. For example, if *S*_*X*_ = 1 then *J* = 1, 1/2, 1/3, 1/4, …, 0, accumulating possible values below 1/2. *S*_*X*_ = 2 then allows for 2/3, which is above 1/2, *S*_*X*_ = 3 allows for 3/4 > 2/3 and so on. Further computation of possible *J* values for increasing *S*_*X*_ (Figure R5) shows an increasingly even distribution of *J*-values. This mechanisms works for any index that can be expressed by Eq. 8 in Šizling et al. (202X), and the effect cannot be eliminated by inventing a new dimensionless index.

**FIGURE R5:**
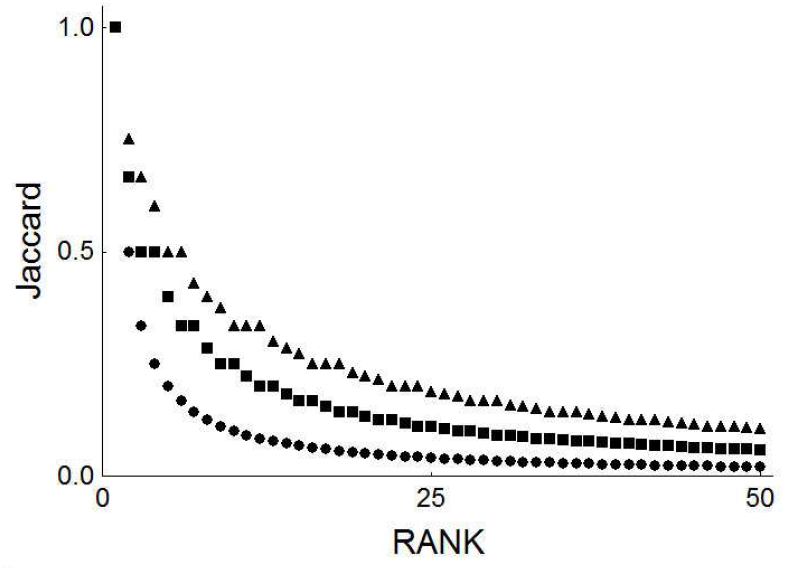
Rank plot of the first fifty values that can reach Jaccard index if *S*_*X*_ is fixed and *S*_*Y*_ and *S*_*X*∩*Y*_ vary within their limits (1 ≤ *S*_*Y*_ < ∞ and 0 ≤ *S*_*X*∩*Y*_ ≤ min(*S*_*X*_, *S*_*Y*_)); the three distributions on display correspond to *S*_*X*_ = 1 (circles), *S*_*X*_ = 2 (squares) and *S*_*X*_ = 3 (triangles).

## R6 (data description)

For comparison with the results from the artificial (simulated) data, we also plotted observed values of the indices. This allows us to identify relationships that are mathematically possible but may be rare or absent in nature. These observations consisted of three different datasets: a set of 29 microbial assemblages extracted from cryoconite on the Greenland Ice Shield, a set of 24 arctic plant assemblages (4 from Greenland, and 20 from Svalbard), and a set of 20 temperate zone plant assemblages (10 from the Czech Republic, and 10 from Southern Norway). The microbial assemblages were sampled by J.Ž. and A.Š, and processed by J.Ž. Plant assemblages were sampled by A.L.Š., Eva Šizlingová and E.T (see Dataset RI.2).

A list of plant species found in a 10x10 m area was recorded. The data are nested in the sense that the assemblages are grouped so that each group of five assemblages is located within a 1km diameter circle. For the purposes of this analysis, only Genera were used.

Microbial assemblages were sampled at 300 m intervals along two lines on the western margin of the Greenland Ice Sheet in the vicinity of Kangerlussuaq. Sampling and sample processing procedures followed Cameron *et al*. 2016. Here we use data inferred from environmental RNA using Illumina amplicon sequencing to detect the active part of the microbial assemblage. Processing of the sequencing output was performed using the QIIME2 environment (Bolyen *et al*., 2019), filtering for sequences present at least three times in the whole dataset, and rarefaction to the sampling depth of 5000 features per sample. This resulted in the exclusion of 7 samples out of 36 that had fewer features than the sampling depth. The remaining samples were used for the diversity analysis.

